# Fate plasticity and reprogramming in genetically distinct populations of *Danio* leucophores

**DOI:** 10.1101/526475

**Authors:** Victor M. Lewis, Lauren M. Saunders, Tracy A. Larson, Emily J Bain, Samantha L. Sturiale, Dvir Gur, Sarwat Chowdhury, Jessica D. Flynn, Michael C. Allen, Dimitri D. Deheyn, Jennifer C. Lee, Julian A. Simon, Jennifer Lippincott-Schwartz, David W. Raible, David M. Parichy

## Abstract

Understanding genetic and cellular bases of adult form remains a fundamental goal at the intersection of developmental and evolutionary biology. The skin pigment cells of vertebrates, derived from embryonic neural crest, are a useful system for elucidating mechanisms of fate specification, pattern formation, and how particular phenotypes impact organismal behavior and ecology. In a survey of *Danio* fishes, including zebrafish *Danio rerio*, we identified two populations of white pigment cells—leucophores—one of which arises by transdifferentiation of adult melanophores and another that develops from a yellow/orange xanthophore-like progenitor. Single-cell transcriptomic, mutational, chemical and ultrastructural analyses of zebrafish leucophores revealed cell-type specific chemical compositions, organelle configurations and genetic requirements. At the organismal level, we identified distinct physiological responses of leucophores during environmental background matching and we show that leucophore complement influences behavior. Together, our studies revealed new, independently arisen pigment cell types and mechanisms of fate acquisition in zebrafish, and illustrate how concerted analyses across hierarchical levels can provide insights into phenotypes and their evolution.

**Significance:** A foundational question in biology is how phenotypically similar traits arise. Here we identify two distinct white pigment cell populations, leucophores, that arise from independent progenitors in zebrafish and its relatives. Remarkably, one of these leucophore populations develops from previously differentiated, black melanophores. These different leucophores exhibited distinct pigment chemistries, cytological features, gene expression profiles and genetic requirements, and whole-animal experiments implicated these cells in behavioral interactions. Our several approaches provide novel insights into pigment cell complements and origins in zebrafish and contribute to our understanding of form and function in the spectacular pigment patterns of teleost fishes.

## Main Text

Vertebrate pigmentation contributes to ecological interactions and is often a target of selection during adaptation and speciation (1–3). Teleost fishes, among the most phenotypically diverse of vertebrate taxa, use a spectacular array of pigment phenotypes to support a variety of behaviors, from attracting mates to predator avoidance, social aggregation, and aggressive interactions. In contrast to birds and mammals that have only a single pigment cell type, the melanocyte, teleosts develop multiple pigment cell classes, including black melanophores, yellow–orange xanthophores, and iridescent iridophores. Pigment patterns reflect the relative abundance and spatial locations of these chromatophores. Decades of work have contributed to understanding developmental and genetic bases of black, yellow and iridescent pigmentation, and cellular interactions underlying pattern formation (4–9).

Teleosts also develop several additional classes of chromatophores, including white or yellow–white pigment cells, or “leucophores” (10). Our knowledge of fate specification, genetic requirements, physical and chemical properties, and behavioral roles of these cells remains fragmentary. Nevertheless, the presumptive origin of all these chromatophores in a common precursor cell population—the embryonic neural crest (11, 12)—and the distinctiveness of these chromatophores in fish that display them, suggest an opportunity to dissect phenotypic diversification ranging from genetic mechanisms and cell fate plasticity to organismal interactions.

Zebrafish *Danio rerio* has emerged as a preeminent laboratory model for studying neural crest development and pigment pattern formation, and comparisons of zebrafish and other *Danio* species have provided insights into the evolution of pattern-forming mechanisms (13–15). Here we investigate physical properties, genetic mechanisms, and cell lineage of leucophores in zebrafish and its relatives. We show that *Danio* fishes have two distinct classes of leucophores with independent developmental origins and cellular architectures. We further identify lineage-specific requirements and pathways modulated in these cells by genetic, chemical and singlecell transcriptomic analyses, and we show that different classes of leucophores can contribute to behaviors at the whole organism level. Our findings suggest that white pigmentation in *Danio* has resulted from phenotypic convergence in neural crest sublineages, providing a novel mechanism for the evolution of teleost pigmentation.

## Results

### Dual Classes and Origins of Leucophores in *Danio*

We evaluated leucophore distribution across 9 species representing multiple subclades within the genus (16). Leucophores containing orange and white pigment were evident in anal fins of 7 species, including zebrafish (in which they had not been reported previously), and less prominently in dorsal fins of 2 species (Fig. 1A,B–left; Fig. S1). Similar to xanthophores, these cells contained pteridines and carotenoids (Fig. S2A–C). During development, yellow/orange pigmentation was evident first, followed by white appearance within ~1 d (Fig. S2D), consistent with genetic analyses in medaka fish that suggest a lineal relationship between leucophores and xanthophores (17, 18). Given the temporal order of pigment deposition in *Danio*, we refer to these cells as xantholeucophores.

**Fig. 1.**
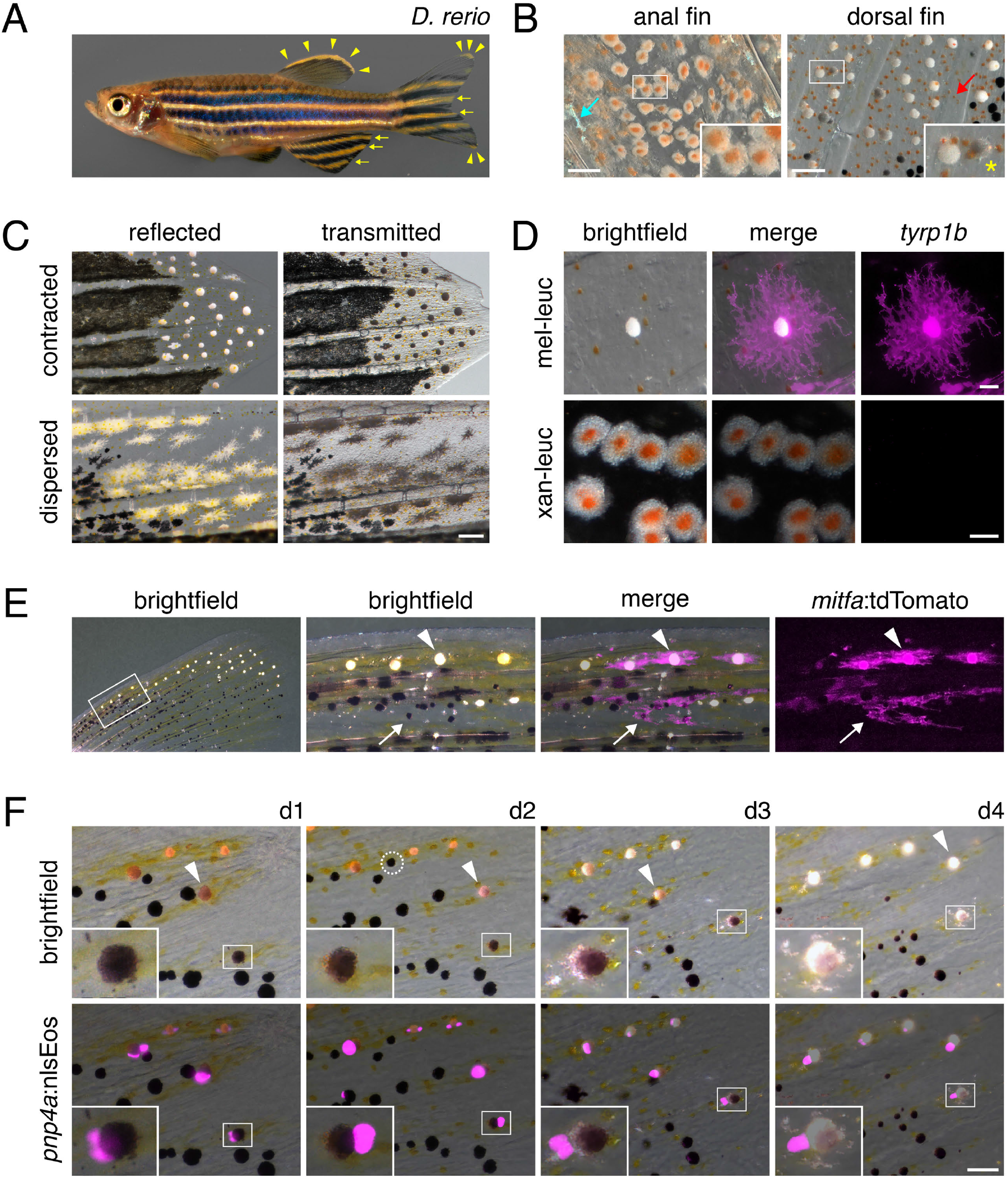
Leucophore appearance and origins in zebrafish. **(A)** Pigment cells containing white pigment. Arrows, leucophores containing yellow/orange pigment. Arrowheads, white cells lacking orange pigment. **(B Left)** Leucophores with white and orange pigment. **(B Right)** Cells containing white pigment. Insets, higher magnification, including grey melanin-containing cell (asterisk). Teal arrow, iridescent iridophore. Red arrow, orange pigment of xanthophore. Fish were treated with epinephrine to contract pigment granules. **(C)** Leucophores at tips of caudal fins under reflected and transmitted illumination. **(D)** Melanoleucophores but not xantholeucophores expressed *tyrp1b*:palm-mCherry. **(E)** Labeled clones contained melanoleucophores (arrowhead) and melanophores (arrow). **(F)** Acquisition of white pigment by a melanophore over 4 days (boxed cell; inset). Arrowhead, a second cell that transits distally. (Scale bars: B, 50 μm; C, 100 μm; D, 20 μm upper, 10 μm lower; E, F, 40 μm)

Besides xantholeucophores, zebrafish has an additional population of leucophores at the distal edges of its dorsal and caudal fins (19). Across species, 8 of 9 *Danio* exhibited leucophores lacking orange coloration at the margin of the dorsal fin, in the dorsal and ventral lobes of the caudal fin, and, in 2 species, within the anal fin (Fig. 1A–B; Fig. S1). These cells had reflective but opaque white material that was variably contracted towards the cell body or dispersed in cellular processes (Fig. 1*C*) and lacked pteridines and carotenoids (Fig. S2*A–B*). Because the cells sometimes contained melanin (Fig. 1B—right; Fig. S1B,C) we refer to them as melanoleucophores.

In zebrafish, melanoleucophores and xantholeucophores expressed distinct pigment cell reporters (Fig. 1D; Fig. S3). Melanoleucophores expressed a *tyrosinase related protein 1b* (*tyrp1b*) transgene that marks melanophores, whereas xantholeucophores expressed an *aldehyde oxidase 5* (*aox5*) transgene that marks xanthophores (12). Both melanoleucophores and xantholeucophores expressed a reporter for *purine nucleoside phosphorylase 4a (pnp4a)*, which is expressed strongly in iridophores and at lower levels in other pigment cell classes (5, 20)

Whereas xantholeucophores resembled leucophores of other species (17, 18), melanoleucophores represent a novel cell type of unknown origin. We hypothesized that melanoleucophores and melanophores share a lineage, given the sporadic melanin found in melanoleucophores and expression by both cell types of the *tyrp1b* transgene. To test this hypothesis, we labeled cells mosaically (21) using an *mitfa:tdTomato* transgene expressed by pigment cell precursors and melanophores (22, 23). Each of 64 labeled clones containing melanoleucophores also included melanophores, indicating that the cells share a common progenitor or that one cell type transdifferentiates from the other (Fig. 1*E*). To distinguish between these possibilities, we labeled pigment cells with photoconvertible Eos (*pnp4a:nlsEos*) and followed cells as they differentiated (12). These analyses demonstrated that melanoleucophores arise directly from melanophores, accumulating white material and losing melanin over several days (Fig. 1F). Consistent with this finding, >95% of newly differentiating melanoleucophores, just beginning to acquire white pigment, contained melanin (143 of 150 cells in 10 larvae). Together these observations indicate that *Danio* develop leucophores of two varieties, with distinct morphologies and developmental origins.

### Distinct Requirements for Specification and Morphogenesis

The origins of xantholeucophores from xanthophore-like cells, and melanoleucophores from melanophores, suggested their development would involve genes required for these respective lineages. Colony stimulating factor-1 receptor-a (Csf1ra) and Kit-a (Kita) promote the development of xanthophores and melanophores, respectively (24–26). Consistent with our prediction, *csf1ra* mutant zebrafish lacked xantholeucophores but retained melanoleucophores, and *kita* mutants lacked melanoleucophores but retained xantholeucophores (Fig. S4A,D). Leucocyte tyrosine kinase (Ltk) is required by iridophores but not xanthophores or melanophores (27), and *ltk* mutants retained both leucophore classes. *D. aesculapii csf1ra* and *kita* mutants exhibited deficiencies identical to those of *D. rerio* mutants (Fig. S4B). Thus, distinct classes of leucophores exhibit distinct genetic requirements.

We asked whether melanoleucophores have genetic requirements different from those of their melanophore progenitors. Because different levels of Kit signaling can promote different cellular outcomes (28, 29), we speculated that melanoleucophores and melanophores might have distinct requirements for Kit activity. We compared phenotypes of fish homozygous for the temperature-sensitive allele, *kita^j1e99^*, at permissive, intermediate and restrictive temperatures (30). While fish reared at restrictive temperatures had neither cell type, fish reared at intermediate temperature developed 91% as many melanophores, but only 14% as many melanoleucophores, as those reared at permissive temperature (Fig. 2A). These findings support a model in which requirements for Kita are greater for melanoleucophores than melanophores.

**Fig. 2.**
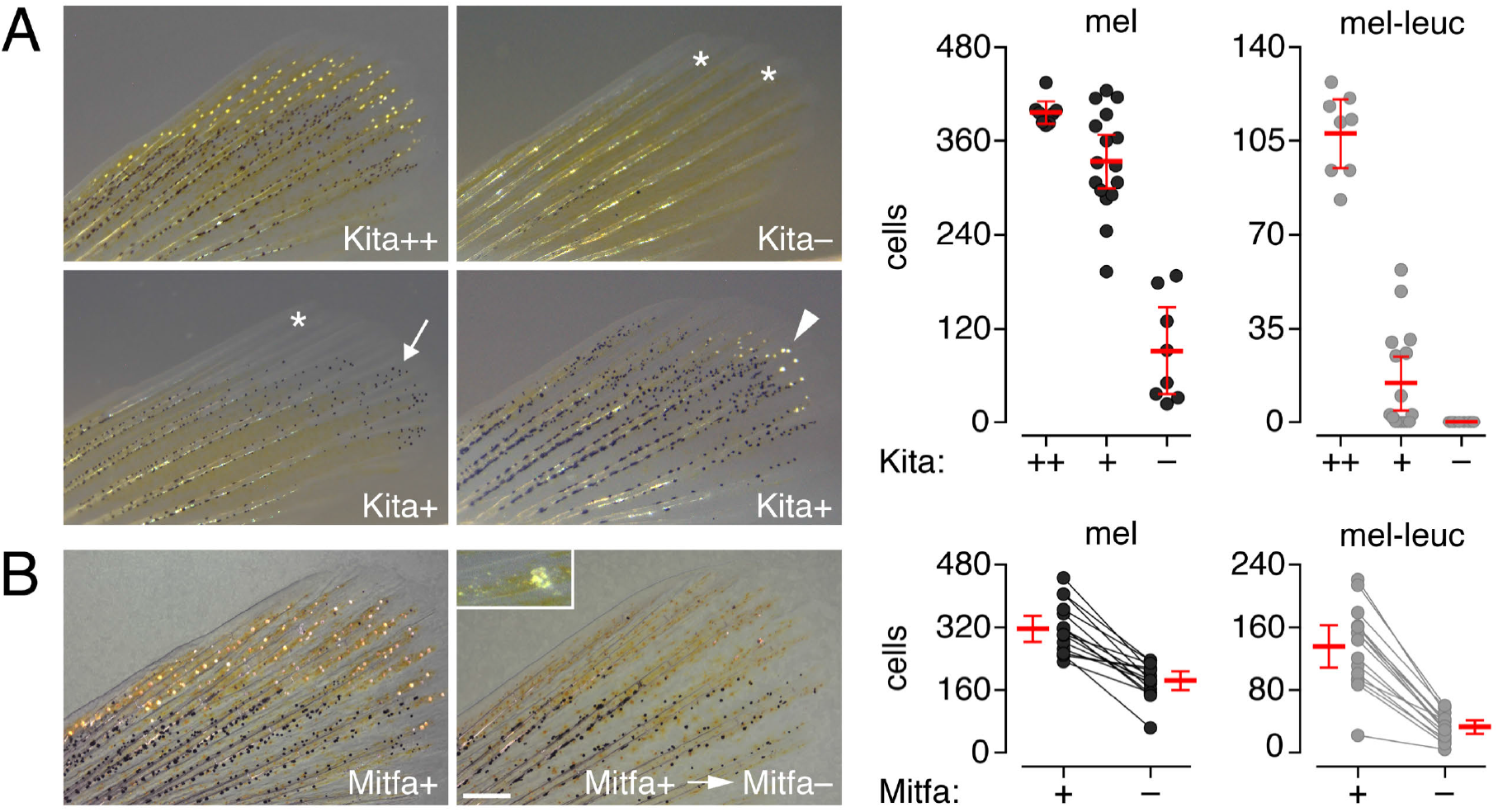
Distinct requirements for Kita and Mitfa in melanoleucophore development. **(A)** *kita^j1e99^* at permissive temperature (Kita++) had wild-type complements of melanophores and melanoleucophores, but at restrictive temperature (Kita–) were deficient for melanophores and lacked all melanoleucophores (asterisks). At an intermediate temperature (Kita+), melanophores were abundant but locations normally harboring melanoleucophores were devoid of cells (asterisk, Left) or populated by melanophores (arrow, Left), or had fewer melanoleucophores compared to permissive temperature (arrowhead, Right). Plots, cell numbers for individual fish (*F*_2,29_ = 65.2 and 119.2, respectively, *P*<0.0001; bars, means ± 95% CI). **(B)** *mitfa^vc7^* fish at permissive temperature (Mitfa+) developed melanophores and melanoleucophores, but many of each died on transition to restrictive temperature (Mitfa–). Panels show same fish before and 2 d after transition. Inset, white-pigment containing cellular debris indicative of cell death [e.g., (59)]. Plots, cell numbers for individual fish before and after transition (paired *t*_15_= −8.6 and −10.9, respectively, *P*<0.0001; bars, means ± 95% CI). Controls maintained at permissive temperature exhibited 1% increases in numbers of each cell type over this period (paired *t*_15_=3.2, *P*<0.05 and paired *t*_15_=2.2, *P*=0.06, respectively). (Scale bar, 250 μm)

The Melanocyte inducing transcription factor, Mitf, is necessary for fate specification, melanogenesis and survival of melanocytes in amniotes and its homologue, Mitfa, has conserved roles in melanophores (31–33). *Danio rerio* and *D. aesculapii* mutants for *mitfa* lacked melanophores and melanoleucophores (Fig. S4C). Unexpectedly, given requirements for Mitfa in melanophore but not xanthophore development (32), we found that *mitfa* mutants also had a xantholeucophore defect, as yellow/orange cells in the position of xantholeucophores lacked white pigmentation (Fig. S4C). To test whether melanoleucophores require Mitfa or whether their deficit reflects the requirement in their melanophore precursors, we employed the temperature-sensitive allele *mitfa^vc7^* (31, 34). We reared fish at permissive temperature to allow melanophore and melanoleucophore differentiation, then shifted fish to restrictive temperature to curtail Mitfa availability. This condition resulted in the loss of 42% of melanophores and 76% of melanoleucophores within 2 d (Fig. 2B). Thus, Mitfa is necessary for at least two stages of the melanoleucophore lineage: development of melanophore progenitors and survival after acquisition of a melanin-free phenotype.

### Distinct Organelle Ultrastructures but Shared Chemistry of Melanoleucophore and Xantholeucophore White Coloration

To address physical and chemical bases for white coloration we examined spectral properties of leucophores. Both melanoleucophores and xantholeucophores reflected across a wide spectrum (Fig. S5A). Xantholeucophores also exhibited a peak of absorption centered at 450 nm even in white regions, indicating the presence of a true pigment. Moreover, both cell types lacked the angular-dependent change of hues—iridescence—of iridophores, which results from stacked arrangements of crystalline guanine reflecting platelets within membrane-bound organelles (35–38) (Fig. 3A). Accordingly, we predicted and then observed that both cell types lack ordered reflecting platelets. Melanoleucophores exhibited variably shaped organelles distinct from reflecting platelets and melanosomes (Fig. *3B,C*). Xantholeucophores were similar to leucophores of other species (35, 39) and xanthophores of zebrafish (38, 40), in harboring round organelles—carotenoid vesicles containing yellow/orange pigment—as well as irregular organelles indistinguishable from xanthophore pterinosomes, which contain yellow or colorless pteridines (Fig. 2D; Fig. S5B).

**Fig. 3.**
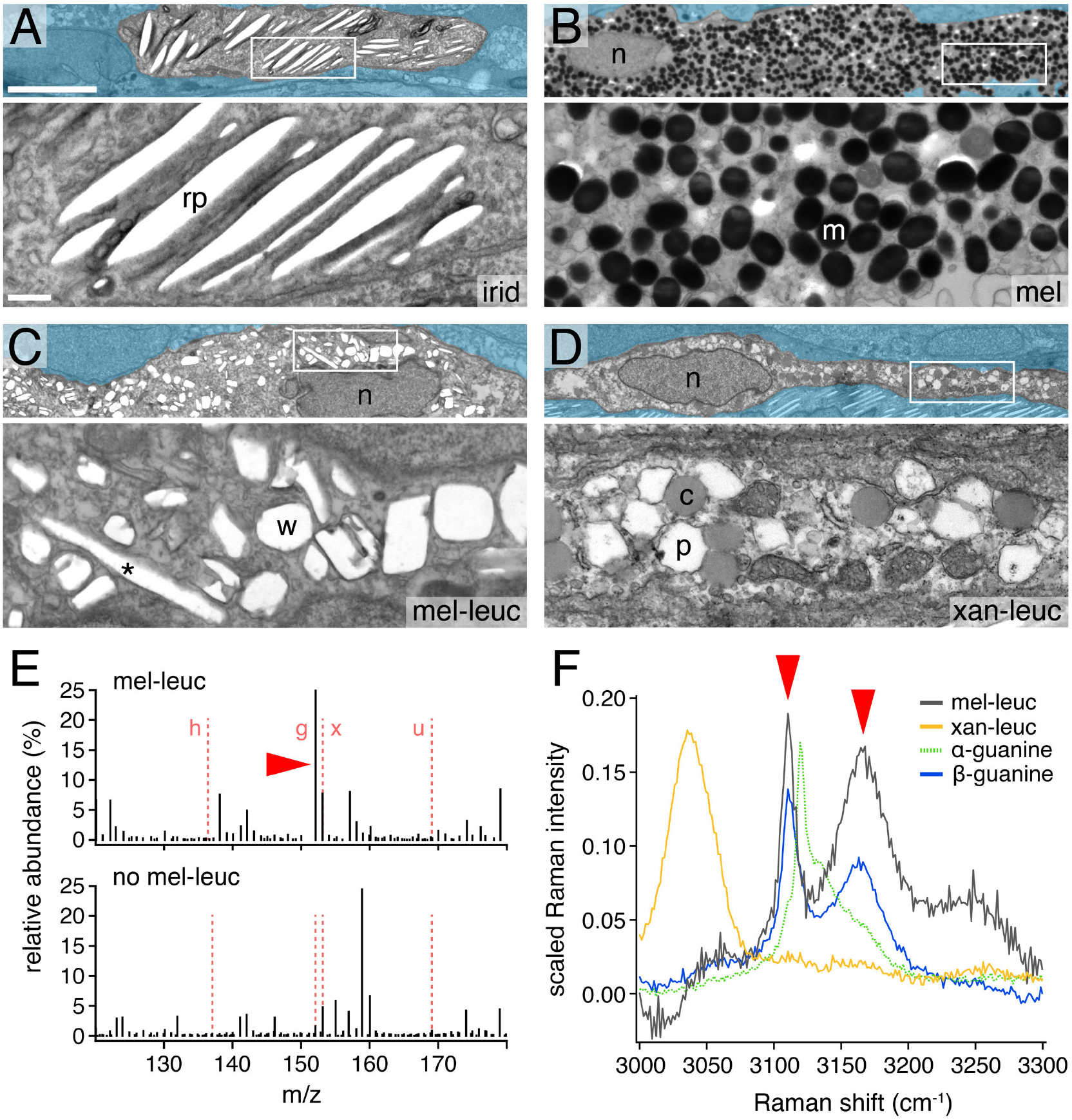
Ultrastructural and chemical characteristics of melanoleucophores and xantholeucophores. **(A)** Iridophore. rp, reflecting platelet. **(B)** melanophore n, nucleus; m, melanosome. **(C)** melanoleucophore. w, presumptive white-pigment containing organelle. **(D)** xantholeucophore. p, pterinosome. Upper panels, low magnification (adjacent cells masked). Lower panels, boxed regions. **(E)** Mass spectrometry of distal fin tissue containing melanoleucophores (upper) revealed more abundant guanine (arrowhead) compared to proximal tissue lacking melanoleucophores (lower; h, hypoxanthine; g, guanine; x, xanthine; u, uric acid). **(F)** Representative Raman spectra of melanoleucophores (grey; *n*=19 cells) and xantholeucophores (orange; n=14) compared to that of α-guanine and β-guanine. High energy peak pattern (arrowheads) indicates β-guanine is present in melanoleucophores but is not detectable in xantholeucophores. (Scale bars, upper 5 μm, lower 500 nm)

We used mass spectrometry to assay purine contents of fin tissue containing different pigment cell classes. Guanine but not other purines was detected in excess in tissues containing melanoleucophores, as compared to fin tissue not containing these cells (Fig. 3*E*; Fig. S5C–D). Raman spectroscopy of individual melanoleucophores indicated the presence of crystalline β-guanine, the meta-stable phase of guanine, typical of iridophores and other biological systems in which crystallization is regulated (41, 42) (Fig. 3F). Tissue harboring xantholeucophores had somewhat increased guanine content (Fig. S5E), though crystalline forms of guanine were not detectable (Fig. 3F), suggesting that colorless pteridines (43, 44) or a combination of factors contribute to white pigmentation. Together, these analyses indicate distinct ultrastructural and chemical bases for white pigmentation of melanoleucophores and xantholeucophores.

### Melanoleucophores and Xantholeucophores Differentially Modulate Pigment Synthesis Pathways

We hypothesized that differentiation of melanoleucophores from melanophores would involve a switch from melanin to purine synthesis and so analyzed transcriptomes of individual pigment cells isolated during melanoleucophore development. Dimensionality reduction and clustering (45) with empirical validation of cluster assignments identified melanophores and melanoleucophores, as well xanthophores and xantholeucophores (Fig. 4A,B; Fig. S6). Supporting a developmental switch in pigmentation pathways, melanoleucophores exhibited higher expression of genes for *de novo* purine synthesis, but lower expression of genes for melanin synthesis, as compared to melanophores (Fig. 4C,D). Pseudotemporal ordering of cells (46) along a differentiation trajectory (Fig. 4B-Right) likewise revealed an inverse relationship in gene expression between *de novo* purine synthesis and melanin synthesis pathways (Fig. 4E,F). This sequence raised the possibility that acquisition of white pigment might depend on prior melanization; yet mutants with unmelanized melanophores developed melanoleucophores, allowing us to reject this model (Fig. S7A).

**Fig. 4.**
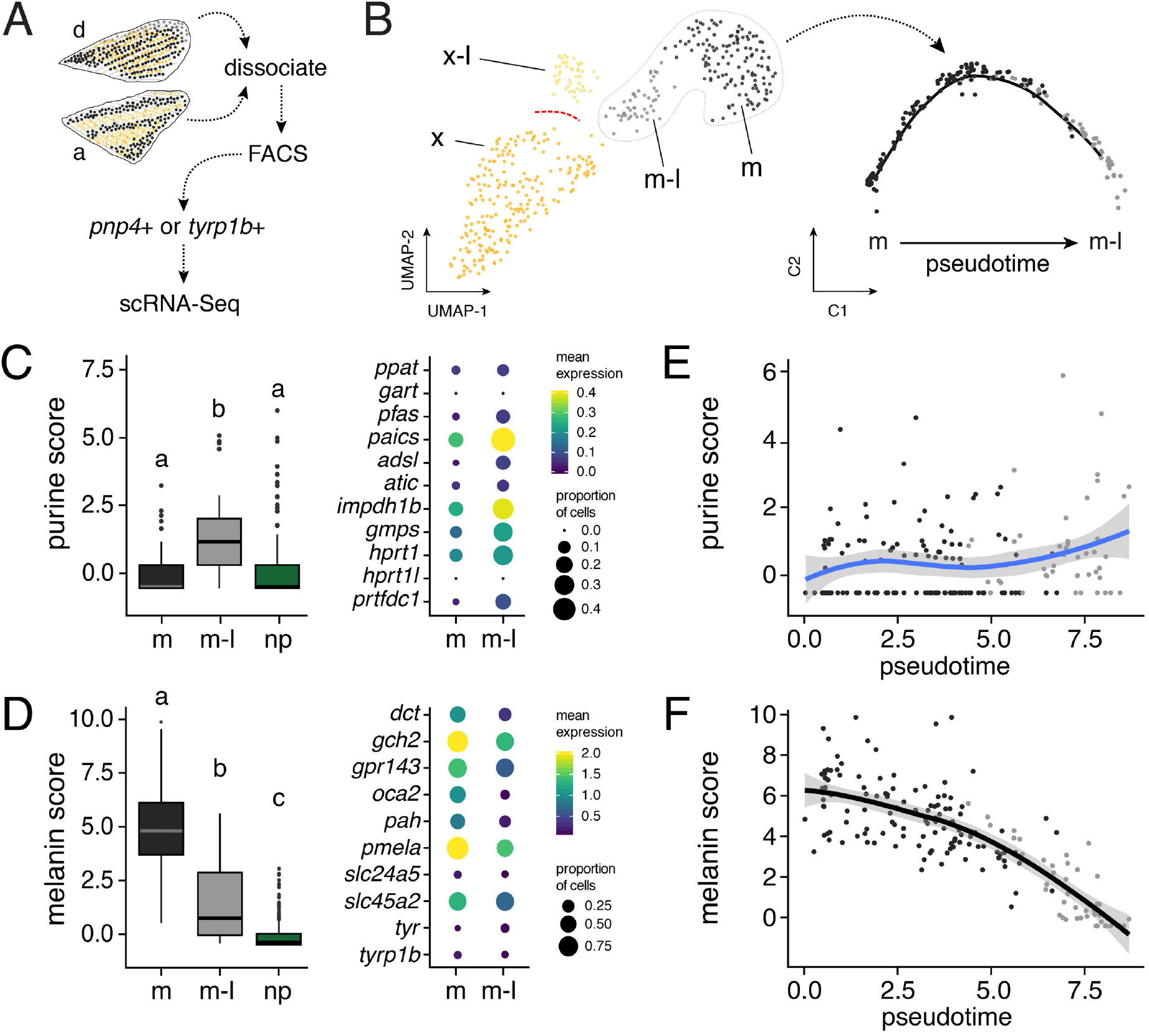
Pigmentation pathway usage and genetic requirements of melanoleucophores. **(A)** Experimental design. d, dorsal fin. a, anal fin. **(B)** Two dimensional UMAP (47) representation of pigment cell clusters (Left; x, xanthophores; x-l, xantholeucophores; m, melanophores; m-l, melanoleucophores) and a differentiation trajectory for melanophores to melanoleucophores (Right). Red dotted line, cell-free region of UMAP space (see main text). **(C)** Aggregate score for de novo purine synthesis genes was greater in melanoleucophores (m-l) than melanophores (m) or non-pigment cells (np). Box plots, medians with interquartile ranges; different letters indicate means significantly different in post hoc comparison (*P*<0.0001; shared letters, *P*=0.2). Dot plots show expression for individual genes. **(D)** Melanin synthesis gene expression was reduced in melanoleucophores compared to melanophores. **(E,F)** Purine and melanin synthesis gene expression scores for individual cells arranged in pseudotime (arbitrary scale) from melanophore (left) to melanoleucophore (right). Line and shaded region, smoothed average with 95% CI.

Compared to melanoleucophores and melanophores, xantholeucophores and xanthophores occupied more distinct locations in transcriptomic space (Fig. 5B-Left), (47). A differentiation trajectory likewise revealed few cells representing intermediate states (Fig. 7B), suggesting that few early differentiating xantholeucophores were recovered from anal fins and that xanthophores (from dorsal fin) are an inadequate proxy for earlier states of xantholeucophore differentiation. Genes involved in purine synthesis, carotenoid processing and pteridine synthesis, were all expressed in xantholeucophores at lower levels than in xanthophores but at higher levels than in non-pigment cells (Fig. S7C,D).

**Fig. 5.**
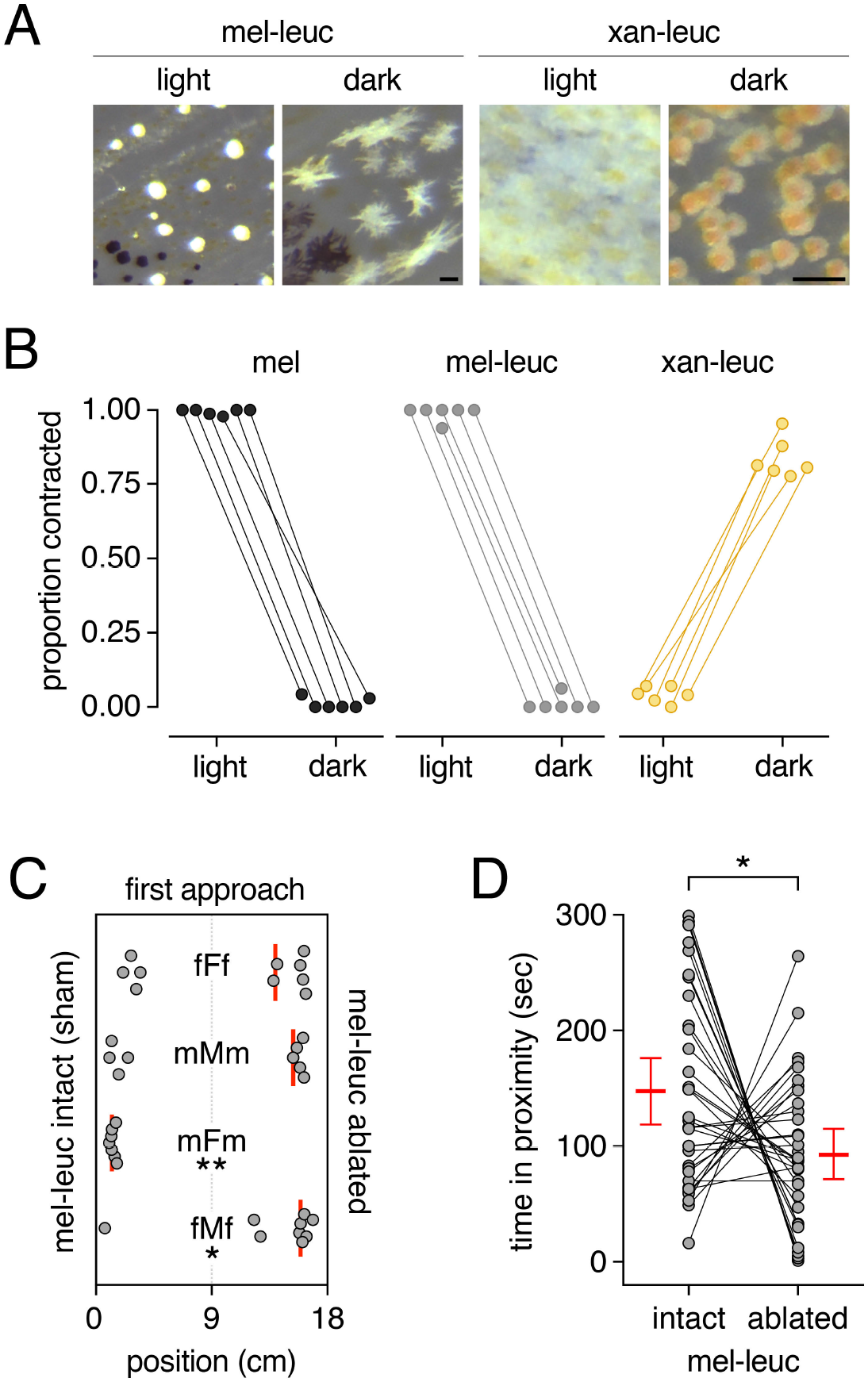
Background adaptation and behavioral responses. **(A)** Cellular morphologies after light and dark background adaptation. **(B)** Melanophores and melanoleucophores exhibited the same responses, whereas xantholeucophores exhibited opposite responses. Points connected by line, cells of a single fish (*n*=26–133 cells, median=62 cells). **(C)** First approach varied with melanoleucophore status and sex. X-axis, horizontal positions in test tank, with test fish released at center and allowed to approach shoals with melanoleucophores intact (sham) or ablated. Experimental conditions: capital letters, sex of test fish; lowercase letters, stimulus shoals of either two males or two females. Each dot represents position reached by a test fish immediately after release. Bars, medians for all test fish. *, X^2^=6.2, *P*<0.05; **, X^2^=9.7, *P*<0.005. **(D)** On average, test fish spent more time in proximity to shoals with intact melanoleucophores than shoals with ablated melanoleucophores (difference in time spent between intact and ablated, null hypothesis mean = 0 sec: two-tailed, *t*_35_=2.29, *P*=0.03). Points connected by lines, times spent by individual fish in proximity to one or another shoal type, with means ± 95% CI. Times did not differ with sex of test fish, sex of stimulus fish, or their interaction (ANOVA, all *P*>0.18). (Scale bars, 20 μm)

### Physiological Responses and Behavioral Implications of *Danio* Leucophores

We sought to understand how leucophores might impact phenotype at the organismal level. During ambient background color adaptation, zebrafish and other species modulate contraction or dispersion of pigment-containing organelles within pigment cells physiologically (38, 39). Typically, melanosomes of melanophores are contracted on a light background, yielding a lighter fish, but are dispersed on a dark background, yielding a darker fish. Leucophores of medaka, containing yellow/orange pigment, have the opposite response of melanophores (48). Given the distinct lineages of melanoleucophores and xantholeucophores, we predicted that their physiological responses to environmental changes would differ. Under standard conditions, pigment of *Danio* melanoleucophores was often contracted whereas that of xantholeucophores was often expanded (Fig. S1B-Top). Melanoleucophores responded, as did melanophores, by contracting pigment in the light and expanding pigment in the dark (Fig. 5A,B). Xantholeucophores displayed the opposite response. Thus, melanoleucophores and xantholeucophores have distinct physiological responses to background adaptation.

Finally, we asked whether social behavior might be influenced by melanoleucophores given their prominent position on the fins, which are held erect during aggressive and courtship interactions (49). To test if zebrafish can perceive and respond to melanoleucophores, we presented fish with alternative shoals having intact melanoleucophores (sham-manipulated) or ablated melanoleucophores (Fig. S8). We assessed which shoal was approached first, and time spent in proximity to one shoal or another (Fig. 5C,D). Test fish with same-sex shoals did not preferentially approach either shoal type. By contrast, test fish with opposite-sex shoals exhibited strong and opposing biases: females were more likely to approach males with intact melanoleucophores; males were more likely to approach females with ablated melanoleucophores. Subsequently, all test fish spent more time in proximity to shoals having intact melanoleucophores. These differences suggest fish attend to the presence of melanoleucophores and raise the possibility of these cells contributing to social interactions in the wild.

## Discussion

It was previously thought that leucophores were most similar to iridophores, however genetic analysis in medaka suggested a shared lineage with xanthophores (7). The phylogenetic distribution of leucophores (Fig. S1), and our finding that melanoleucophores transdifferentiate from melanophores while xantholeucophores differentiate from xanthophore-like progenitors, suggest that a *Danio* ancestor evolved two types of leucophores independently (Fig. S9A). The finding that white bars of clownfish *Amphiprion* depend on iridophores (50) show that each of the three major chromatophore lineages can contribute to a white pigmentary phenotype.

An unresolved question in the evolution of morphology is the extent to which similar but independently evolved phenotypes depend on the same (parallel) or different (convergent) mechanisms. The deposition of guanine crystals in iridophores and melanoleucophores suggests a parallel cooption of purine synthesis and biomineralization, whereas xantholeucophores appear to depend on a convergent solution to achieving a white pigmentary phenotype. Moreover, each of these cells has a unique organelle morphology and arrangement (Fig. S9B), indicating convergence at the level of cellular architecture. These observations illustrate the value of concerted analyses at multiple hierarchical levels in dissecting trait origins.

Our findings also provide insights into how known pigmentary genes have been utilized in the evolution of white coloration. The dependence of xantholeucophores and melanoleucophores on xanthogenic *csf1ra* and melanogenic *kita*, respectively, supports a model in which leucophore deployment is constrained by prior lineage requirements (Fig. S9A). Our analysis revealed a novel role for *mitfa* in xantholeucophore development and conditional genetic analyses revealed roles for *kita* and *mitfa* specific to melanoleucophore development. It remains for future studies to interrogate how these and other “classic” pigmentation genes integrate with pathways required for the white pigmentary phenotype—including upregulation of *de novo* purine synthesis, downregulation of melanin synthesis, and factors underlying melanin depletion. An intriguing possibility is that *mitfa* itself plays a role, analogous to functions in fate specification or phenotype switching in melanocytes, melanophores and melanoma (51).

Finally, white pigmentation is thought to be ecologically relevant: white bars of clownfish contribute to species recognition (52), a yellow-white bar in the caudal fin of male Goodeinae fishes is linked to mating success (53, 54), and white pigmentation in fins of guppy, *Poecilia reticulata*, likely enhances honest signaling in sexual selection (55). Our work reveals contributions of xantholeucophores and melanoleucophores to background adaptation, and potential roles for melanoleucophores in social interactions, especially given the anatomical positions of these cells, and their likely visibility in the variable habitats experienced by zebrafish and other *Danio* in the wild (13, 56).

## Materials and Methods

### Fish Rearing, Lineage Analysis and Temperature Shift Experiments

Zebrafish and other *Danio* species were housed at ~28 °C, 14L:10D, fed rotifers, *Artemia* and flake food. Clonal labeling and Eos-fate mapping followed (12, 21). Fish were anesthetized in MS222 prior to imaging or fin clipping. Protocols were approved by Institutional Animal Care and Use Committees of University of Virginia and University of Washington.

### Physical and Chemical Analyses

Measurement of reflectance and transmittance used a PARISS hyperspectral imaging system mounted on a Nikon Eclipse 80i microscope. TEM used standard methods. Guanine content was analyzed on an Agilent HPLC with photodiode array detector and single quadrupole mass spectrometer after extracting purines in 1M NaOH, and using a custom-built Raman microscope.

### Single-Cell RNA-Sequencing and Analysis

Cells expressing tyrp1b:palm-mCherry or pnp4a:palm-mCherry were isolated by fluorescence activated cell sorting, captured in a Chromium controller (10X Genomics) and sequenced on an Illumina NextSeq 500. Visualization used UMAP (47) with clusters validated using cells of specific phenotypes collected manually and tested by RT-PCR. Trajectory analysis used Monocle (v2.99.1). Gene sets for signature scores were manually curated from the literature and ZFIN.

### Physiological and Behavioral Response Testing

Fish were tested for background adaptation in black or white beakers under constant light for 5 minutes, with scoring after (35). Behavioral assays (57, 58) tested for effects of sham-mamipulation or melanoleucophore ablation as well as sex of both test and stimulus fish.

## Acknowledgements

We thank J. Liu for observations and discussions related to xantholeucophores, A. Schwindling and D. White for assistance with fish rearing, A. Wills for microscope use and E. Parker for assistance with TEM. Supported by NIH R35 GM122471 to DMP, NEI P30 EY001730 Core Grant for Vision Research to UW Department of Ophthalmology Vision Core, MURI-Melanin AFOSRT FA9550-18-1-0142 to DDD. JDF and JCL were supported by the Intramural Research Program of the National Institutes of Health (NIH), National Heart, Lung, and Blood Institute.

## Supplementary Information for

**Supplementary Information Text**

Figures S1–S9 Materials and Methods Tables

**Fig. S1.**
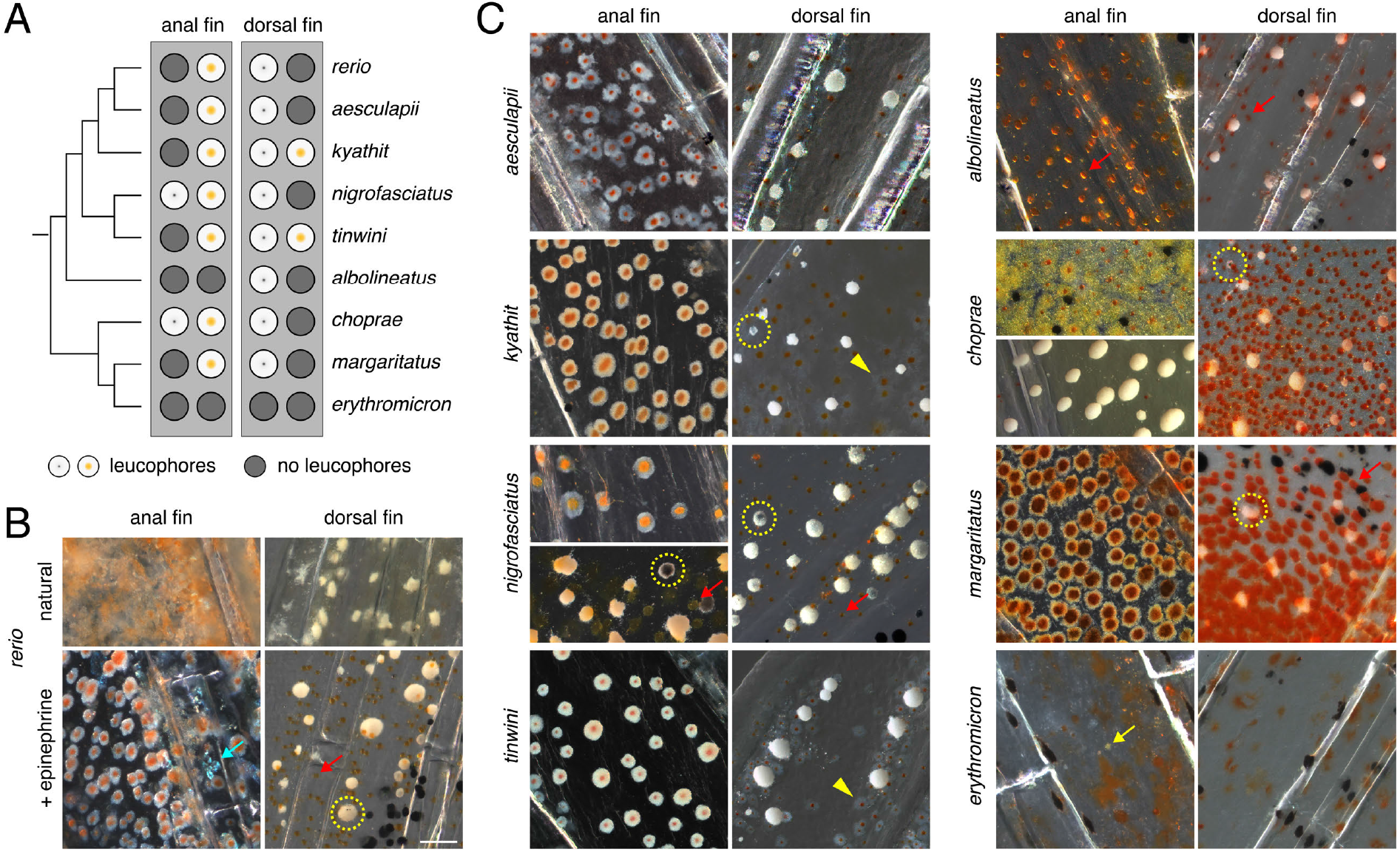
Widespread distribution of leucophores in *Danio*. **(A)** Phylogenetic relationships of select *Danio* [after (1)] indicating presence in anal and dorsal fins of leucophores either containing white and orange pigment (“xantholeucophores”) or white and occasional black pigment (“melanoleucophores,” see Main Text). Only *D. erythromicron* lacked such cells. For images of whole fish see (2). **(B)** Examples of xantholeucophores in the anal fin and melanoleucophores in the dorsal fin of *D. rerio*. Upper panels show morphologies of cells in fish housed in standard rearing conditions. Lower panels show cellular phenotypes following treatment with epinephrine. Teal arrow, an iridescent iridophore. Red arrow, contracted orange pigment of xanthophore, lacking discernible white pigment. Dotted circle, melanoleucophore containing melanin. **(C)** Examples of xantholeuocophores and melanoleucophores, or their absence, in fins of other *Danio* shown in *A*. Note that both leucophore classes were evident in anal fins of *D. nigrofasciatus* and *D. choprae*, with melanoleucophores located distally and xantholeucophores more proximally. The orange or yellow cast to anal fin melanoleucophores in *D. nigrofasciatus* and *D. choprae* results from co-occuring xanthophores, individual outlines of which are not apparent in these views. Yellow arrowheads, less distinctive xantholeucophores in dorsal fins of *D. kyathit* and *D. tinwini*. Red arrows, orange or red pigment of xanthophores or erythrophores, respectively. Yellow arrow, white debris of unknown origin in anal fin of *D. erythromicron*. Other annotations as in B. (Scale bar in B for B and C, 50 μm)

**Fig. S2.**
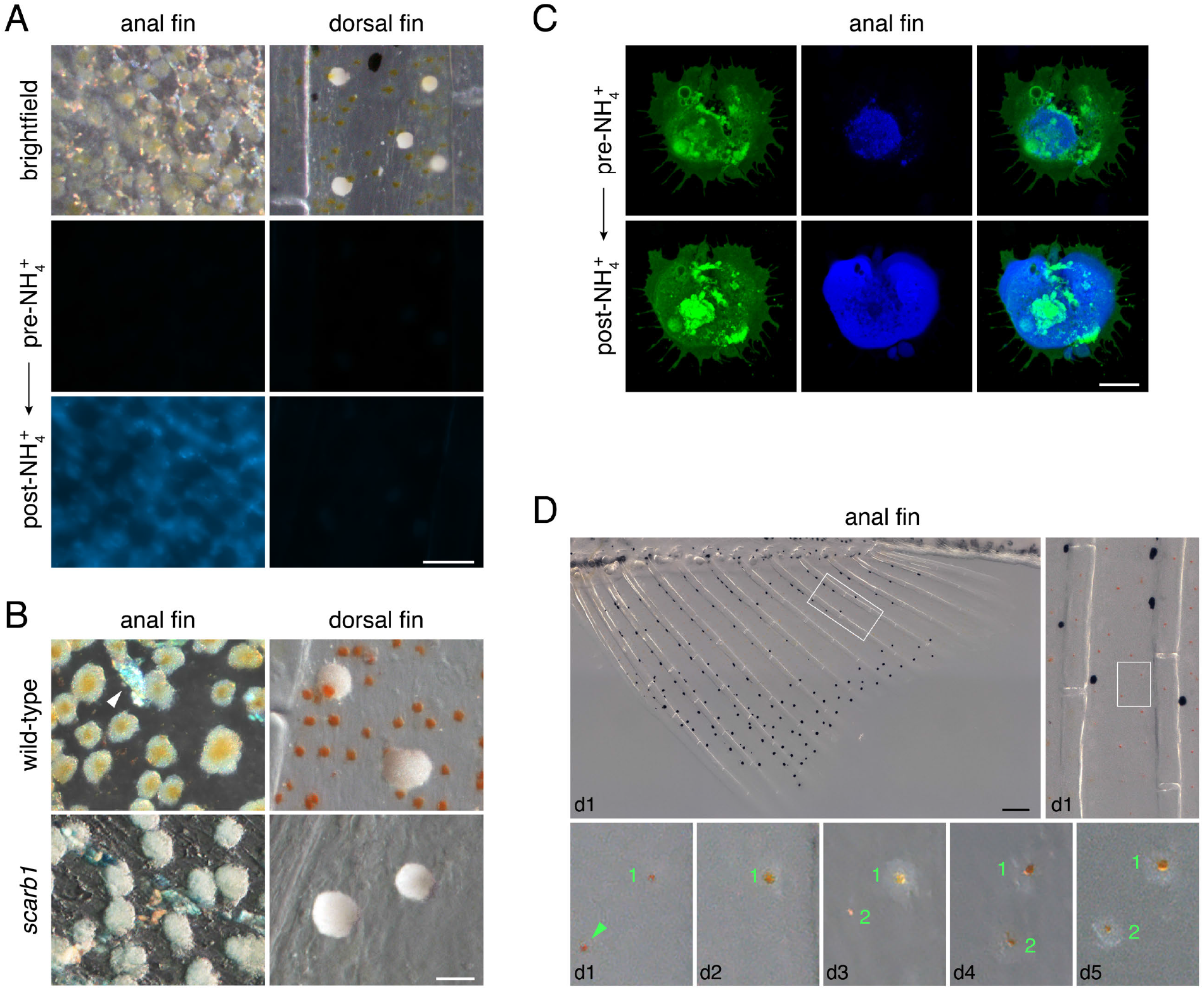
Pigmentary components of xantholeucophores and first appearance. **(A)** Similar to xanthophores, leucophores of the anal fin (xantholeucophores) contained pteridines (3), revealed by autofluorescence under DAPI illumination in response to dilute ammonia treatment (4). Before administration of ammonia (pre-NH_4^+^_), no fluorescence was detectable at this magnification and exposure. After ammonia treatment (post-NH_4^+^_), xantholeucophores exhibited strong autofluorescence whereas leucophores of the dorsal fin (melanoleucophores) did not exhibit autofluorescence. Image exposure times were the same before and after treatment and between cell types. **(B)** Yellow-orange coloration of anal fin xantholeucophores is contained pteridines is carotenoid-based, as indicated by the absence of such color in zebrafish mutants for *scarb1*, an essential gene for carotenoid localization (5). Dorsal fin melanoleucophores were indistinguishable between wild-type and *scarb1* mutant fish. Arrowhead, iridophore. **(C)** High resolution image of live, unmanipulated xantholeucophore plated *ex vivo*, reveals carotenoid vesicles and background autofluorescence (green) with weak autofluorescence centrally in DAPI channel (blue). Treatment of the same cell with dilute ammonia induced widespread autofluorescence (blue) indicative of pteridine release. (D) Anal fin xantholeucophore first appearance in *D. aesculapii* [equivalent to PR stage (6) *D. rerio*]. Images on day 1 (d1) indicate two representative, orange-pigment containing xanthophore-like cells, and their broader tissue context. One cell (arrowhead) migrated from the field of view whereas another, cell (1) began to acquire white pigment on d2, which was evident more prominently by d3. A second newly appearing cell (2) on d3, began to acquire white pigment by d4 and had clearly evident white pigment by d5. (Scale bars: A, 50 μm; B, 20 μm; C, 10 μm; D, 200 μm)

**Fig. S3.**
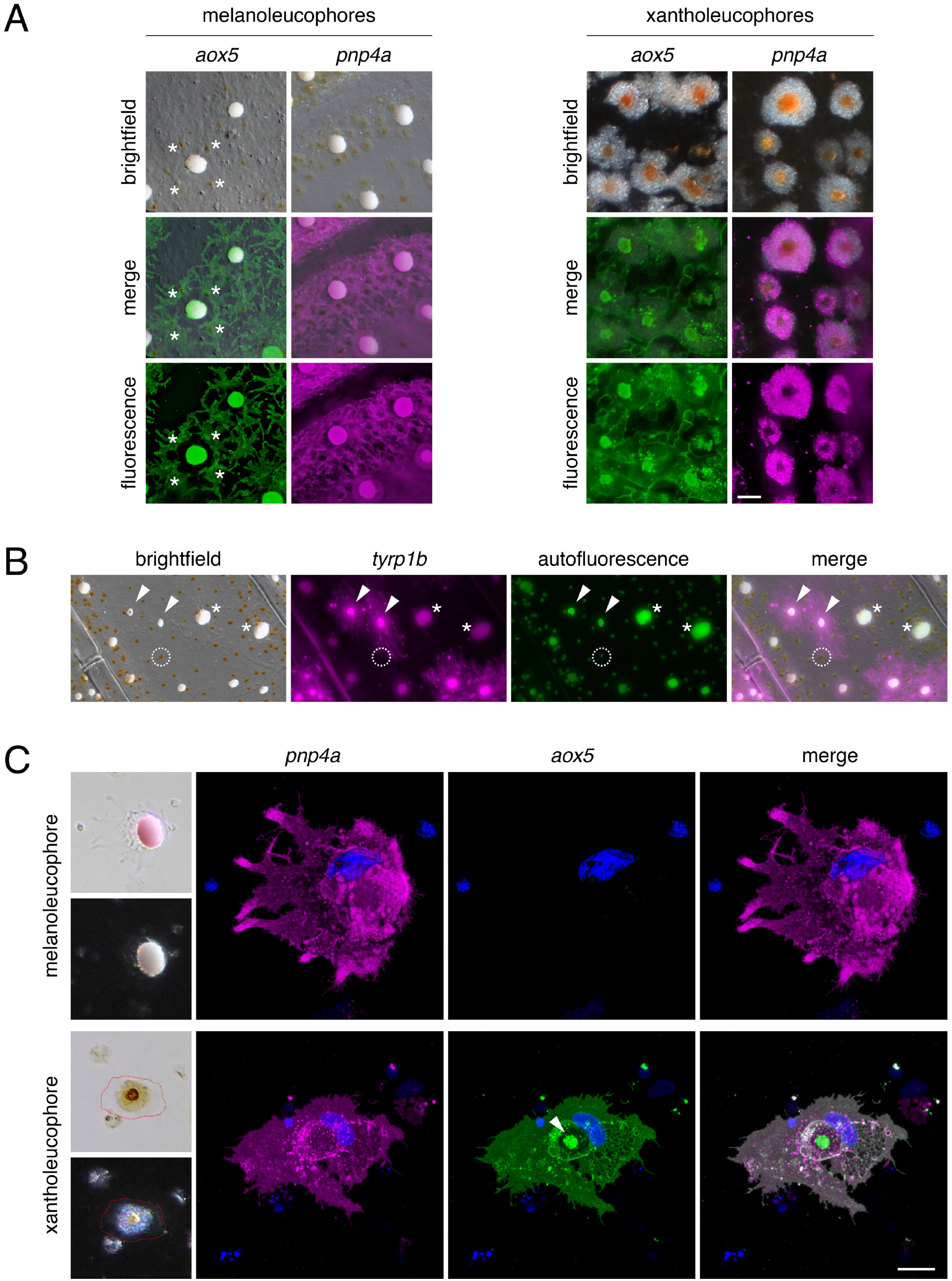
Reporter labeling. **(A)** Brightfield and fluorescence images for aox5:palm-EGFP and *pnp4a*:palm-mCherry. For dorsal fin, asterisks indicate *aox5* transgene expression by xanthophores adjacent to a melanoleucophore. **(B)** Mosaic expression of *tyrp1b* transgene in melanoleucophores. Arrowheads, two cells that expressed the transgene as evidenced by membrane-targeted palm-mCherry expression. Cells with small amounts of white pigment and some melanin often expressed *tyrp1b*:palm-mCherry (arrowheads), whereas cells having larger accumulations of white pigment and no melanin (asterisks) failed to express detectable levels of reporter. Punctate fluorescence centrally was reflectance from white pigment, evident also in GFP channel; dashed circle indicates autofluorecence of carotenoids in xanthophores. **(C)** High resolution images of live melanoleucophore and xantholeucophore plated *ev vivo* illustrate expression of *pnp4a* transgene by both cell types but xantholeucophore-specific expression of *aox5*. Arrowhead, compacted, autofluorescing carotenoids (contained within carotenoid vesicles) contracted to cell center by epinephrine. Blue, DAPI-labeled nuclei. Note that some cell shape change is evident between bright-field and fluorescence images, which were acquired on different microscopes. Approximate boundaries of xantholeucophore are outlined for clarity in brightfield. (Scale bars: A, 20 μm; B, μm; C, 10 μm)

**Fig. S4.**
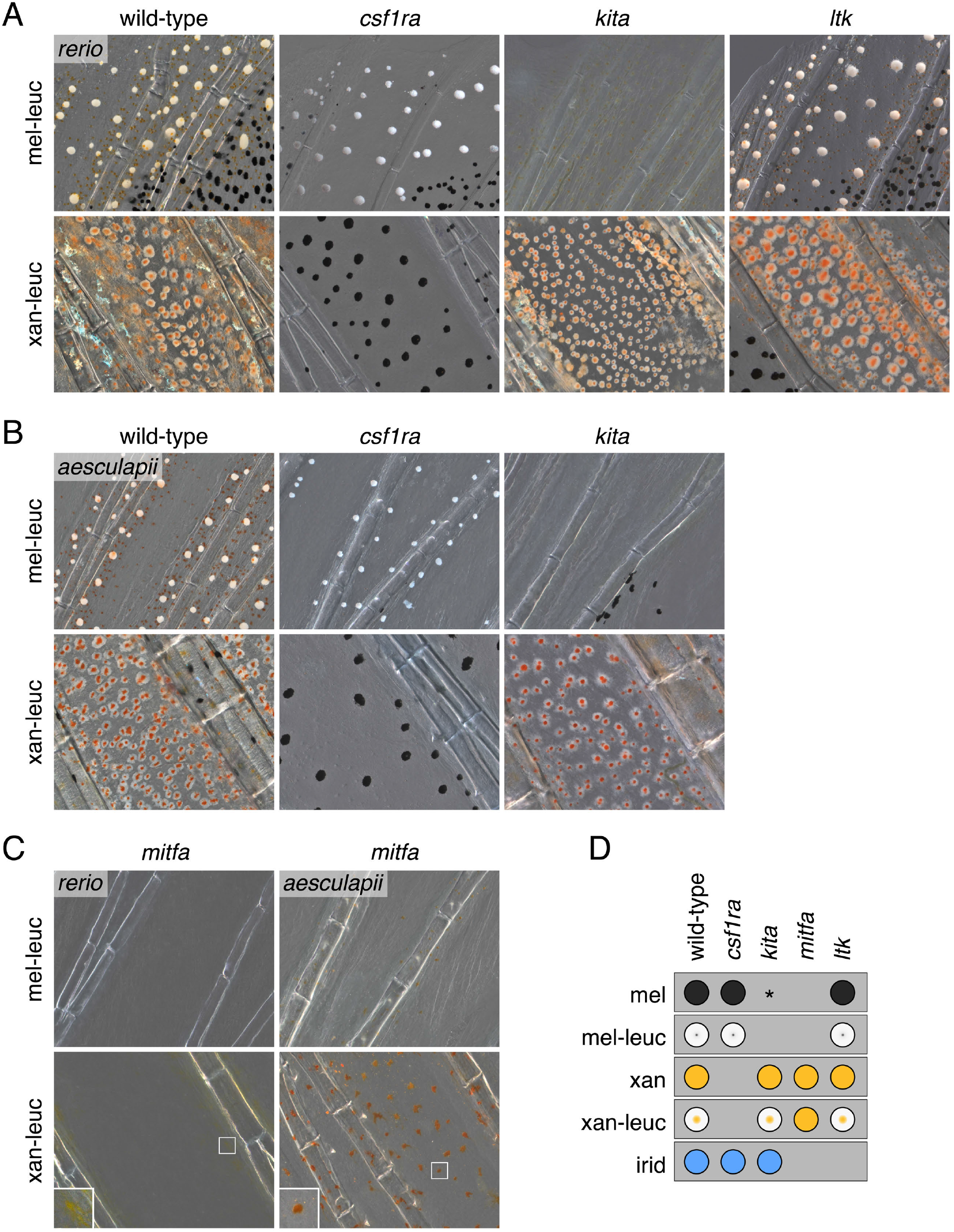
Genetic requirements of melanoleucophores and xantholeucophores. **(A)** In *D. rerio*, fish mutant for a presumptive null allele of *csf1ra* (7) retained melanoleucophores, while lacking xanthophores. By contrast, melanoleucophores were absent from fish mutant for a null allele of *kita*, in which melanophores are deficient (8). Mutants exhibited reciprocal phenotypes for xantholeucophores, which were lost in the *csf1ra* mutant, but persisted in the *kita* mutant. An *ltk* mutant, completely lacking iridophores (9), retained both leucophore classes. **(B)** *D. aesculapii* mutant phenotypes were concordant with those of *D. rerio*. It was not feasible to test genetic requirements of leucophores for *ltk* in *D. aesculapii* owing to lethality of null alleles (9). **(C)** In mutants homozygous for presumptive null alles of *mitfa* (10), melanoleucophores were lost in addition to melanophores. White-pigmented xantholeucophores as well as iridophores were lost from fins, despite persistence of xanthophores and iridophores elsewhere on the fish (10). Insets, cells with yellow or orange pigment that may be xanthophores or white-pigment free xantholeucophores. **(D)** Summary of genetic analyses. Asterisk, melanophores occur sparsely in proximal regions of caudal fin and body of *D. rerio kita* mutants (8, 11).

**Fig. S5.**
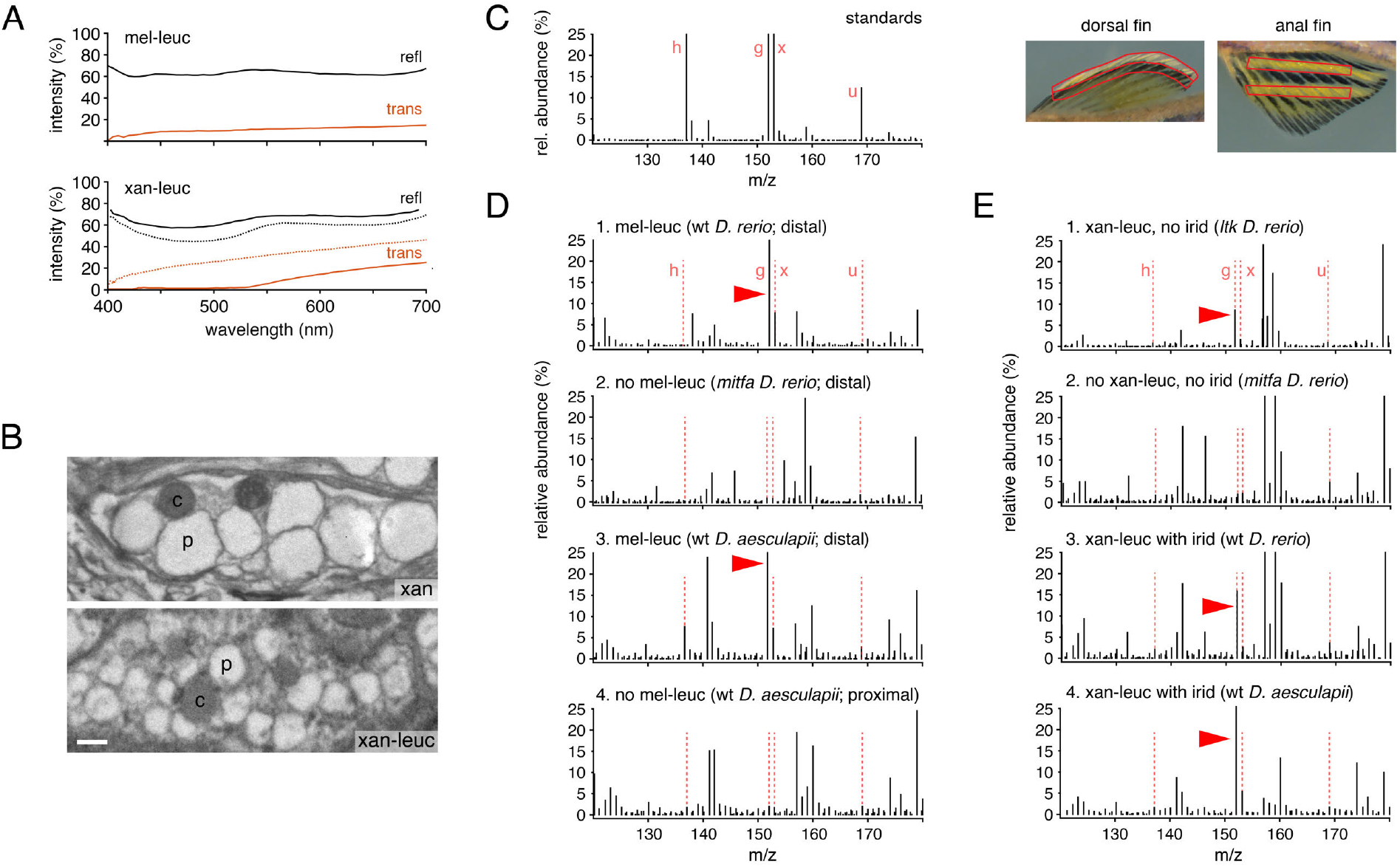
Physical and chemical characteristics of melanoleucophores and xantholeucophores. **(A Upper)** Hyperspectral imaging of melanoleucophores revealed high, nearly uniform reflectance (refl) across visible wavelengths and correspondingly low transmittance (trans). **(A Lower)** Hyperspectral imaging of xantholeucophores was performed across regions devoid of carotenoid (solid lines) and regions in which carotenoids had been contracted to cell centers by epinephrine (dotted lines). Reduced reflectance and and transmittance below ~550 nm are consistent with the presence of pteridines and carotenoids not found in melanoleucophores. **(B)** Xanthophores and xantholeucophores both contained pterinosomes (p) and carotenoid vesicles (c). Organelle sizes were variable within and between cell types. **(C)** Mass spectrometry standards (h, hypoxanthine; g, guanine; x, xanthine; u, uridine) and fin regions excised for mass spectrometry. In dorsal fin, tissues with and without melanoleucophores were compared. From anal fin tissue, regions containing xantholeucophores (and some iridophores) were compared between genetic backgrounds with or without these cells. **(D)** Supplementary mass spectrometric analysis of dorsal fin tissue with and without melanoleucophores. Plots *D-1* and *D-2* indicate illustrate guanine peak in distal fin tissue containing melanoleucophores of wild-type *D. rerio* (same as Fig. 2E—upper) and loss of this peak from the same region of fin when melanoleucophores are ablated genetically in the *mitfa* mutant (Fig. S4C). D-3 and D-4 illustrate co-occurrence of excess guanine with melanoleucophores in *D. aesculapii*, as in *D. rerio* (compare to Fig. 3E). **(E)** Mass spectrometric analysis of anal fin tissue. Plot E-1 shows detectable guanine in fin tissue containing xantholeucophores but not iridophores of *ltk* mutant *D. rerio* (Fig. S4A). E-2, Guanine peak is lost when both xantholeucophores and iridophores are lost in *mitfa* mutant *D. rerio*. E-3, Increased guanine peak of wild-type *D. rerio* fin containing both xantholeucophores and iridophores. E-4, xantholeucophore- and iridophore-containing fin of *D. aesculapii* has excess guanine similar to wild-type *D. rerio*. (Scale bar in B, 500 nm)

**Fig. S6.**
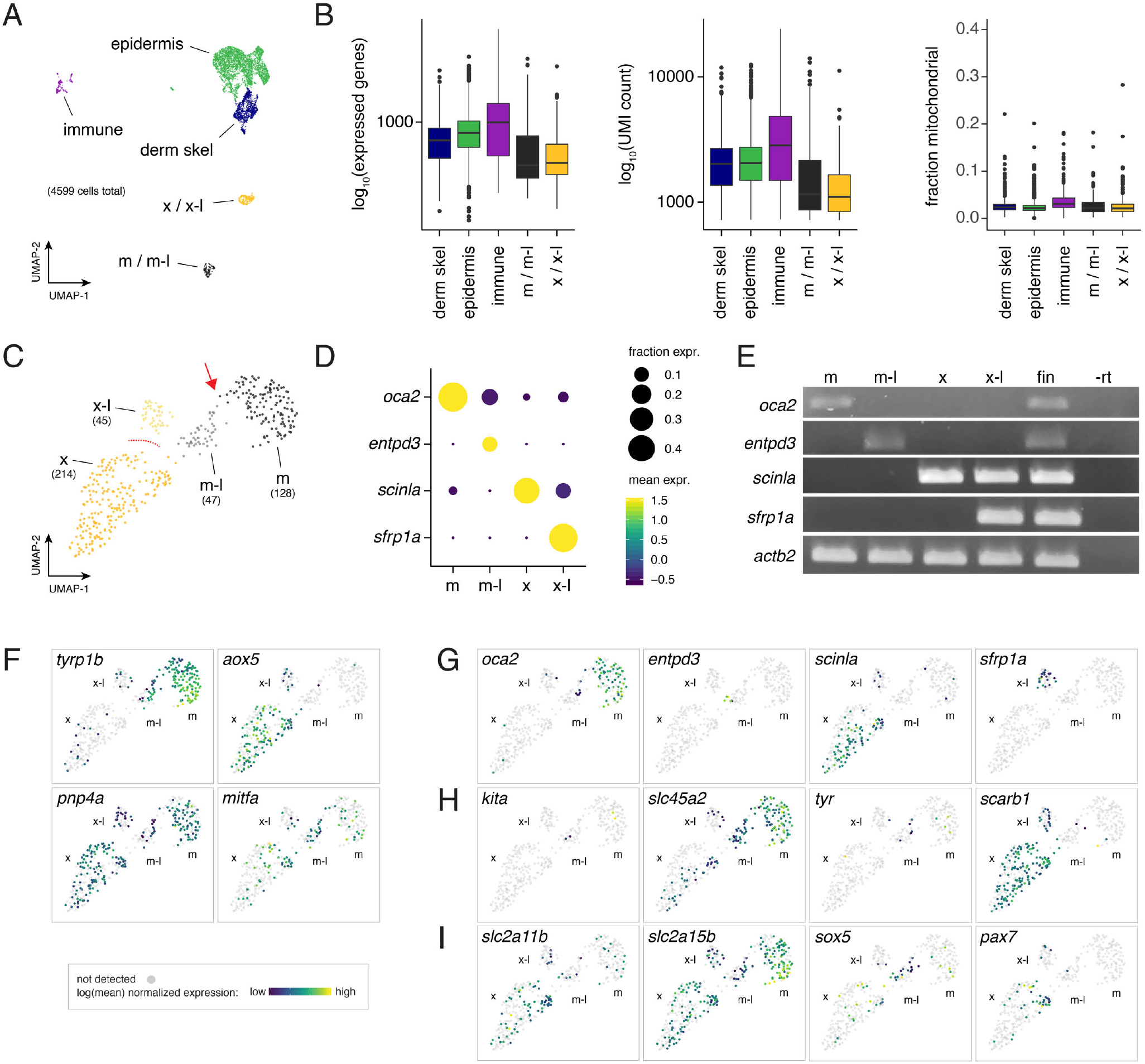
Population characteristics and clustering of single cell RNA-sequencing (scRNA-Seq) data set from zebrafish fin. **(A)** UMAP representation of all cells collected (*N*=4599) with identities of major clusters inferred by expression of known markers, including cells of epidermis, dermal skeleton, and the immune system as well as xanthophores or xantholeucophores (x/x-l) and melanophores or melanoleucophores (m/m-l). Only small numbers of iridophores occur in fins of *Danio* fishes and clusters of these fragile cells were not recovered in these analyses. **(B)** Counts of unique expressed genes, unique molecular identifiers (UMI) and fractions of mitochondrial reads by cell type. Relatively low proportions of mitochondrial genes observed here are consistent with intact rather than broken cells (12). Plots show medians with boxes spanning interquartile ranges; vertical lines indicate farthest observations with outliers depicted individually. **(C)** UMAP representation of pigment cell clusters inferred by molecular marker expression to represent xanthophores (x), xantholeucophores (x-l), melanophores (m) and melanoleucophores (m-l). (Same plot as Fig. 4B-left.) Numbers in parenthesis indicate total cells comprising each cluster. Arrow indicates cells bridging melanophore and melanoleucophore clusters. Dashed lines indicate lack of cells occupying space between xanthophores (derived from dorsal fin) and xantholeucophores (derived from anal fin). **(D,E)** *Post hoc* verification of cluster cell-type assignments by comparison of visually confirmed cellular phenotypes and gene expression. Plot in D illustrates mean expression level and proportion of cells expressing each of 4 candidate marker loci across the 4 pigment cell clusters identified by scRNA-Seq and UMAP clustering. RT-PCR in *E* illustrates expression of same marker loci in cells of each class picked manually *in vitro*, as well as whole fin tissue (RT–, fin tissue prepared without reverse transcriptase). (F–I) Gene expression mapped to UMAP representation of cell clusters. **(F)** Genes corresponding to reporters used in this study. Breadth of *pnp4a* and *mitfa* expression were consistent with independent observations (5). **(G)** Genes used in *post hoc* validation of cell-type assignments (D,E). **(H)** Genes corresponding to mutants used in this study *(csf1ra* expression was not detectable by scRNA-Seq). **(I)** Genes identified for roles in ealrly larval leucophore development in medaka. *slc2a11b* corresponds to medaka *white leucophore* (*wl*), required for the development of yellow coloration in both xanthophores and leucophores. *slc2a15b* (*leucophore free, lf*), is required for both leucophore and xanthophore development from early stages. *sox5* (*many leucophores, ml-3*) promotes the development of xanthophores over leucophores. *pax7a* (*lf-2*) is required for the development of most leucophores and all xanthophores. None of the medaka leucophore-associated genes exhibited strongly type-specific expression in zebrafish adult fin pigment cells.

**Fig. S7.**
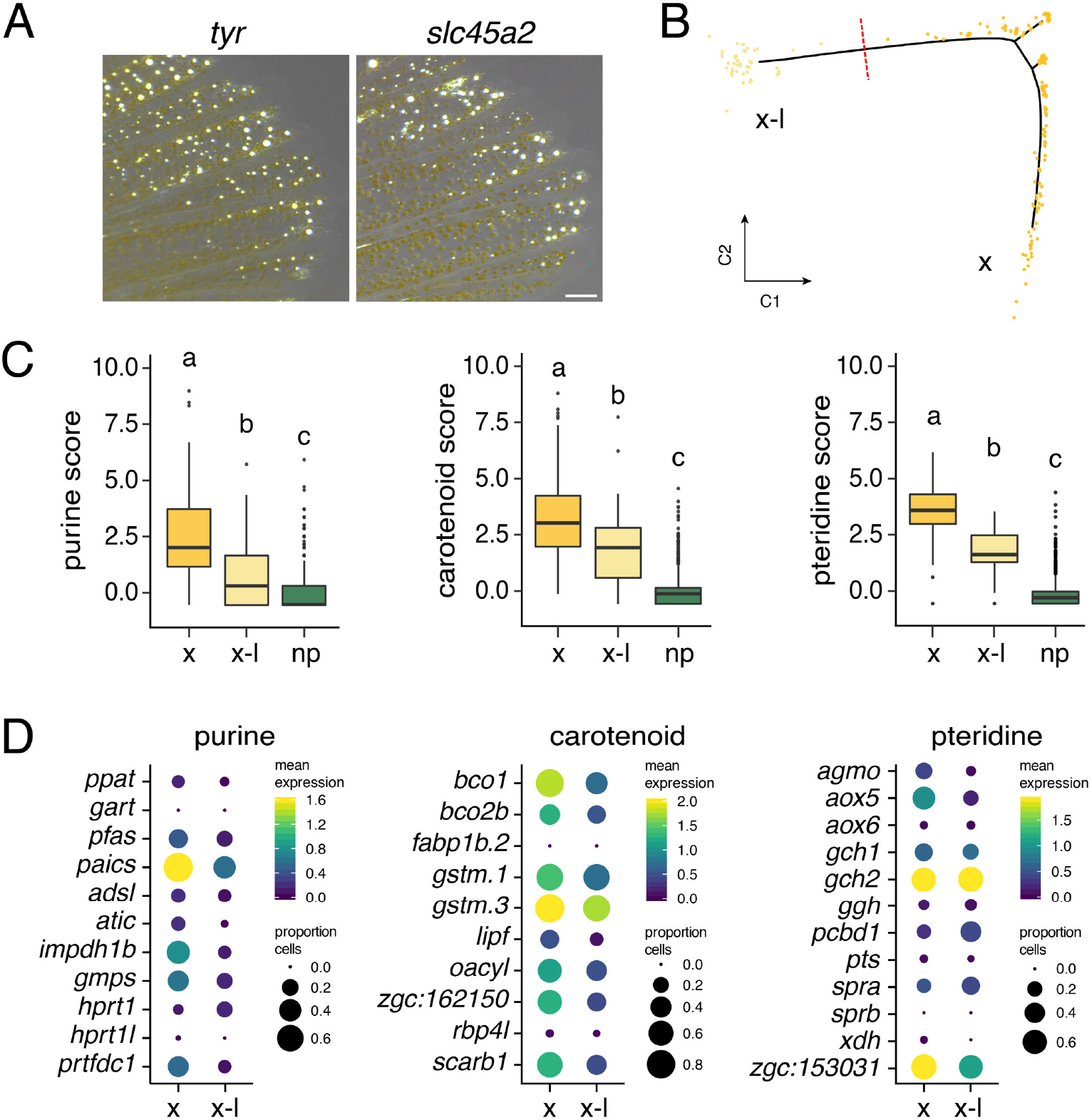
scRNA-Seq and genetic pathway analysis of melanoleucophore differentiation and xantholeucophores relative to xanthophores. **(A)** Melanin-free *D. rerio* mutants of *tyrosinase* (*tyr*) and *albino* (*slc45a2*) retain normal leucophores. **(B)** Pseudotemporal ordering of xanthophores (x) and xantholeucophores (x-l) revealed few cells where an intermediate state might be found (red dotted line). Most xanthophores were collected from dorsal fin whereas xantholeucophores were collected from anal fin. **(C,D)** Aggregate expression scores (C) and individual expression profiles (D) for genes of purine, carotenoid and pteridine pathways in xanthophores and xantholeucophores compared to non-pigment cells (np). Different letters above bars indicate means significantly different in *post hoc* comparisons (all *P*<0.001). Guanosine triphosphate is the most basal substrate for pteridine synthesis (3) so higher expression of purine synthesis genes by xanthophores and xantholeucophores compared to non-pigment cells is expected. (Scale bar, 250 μm)

**Fig. S8.**
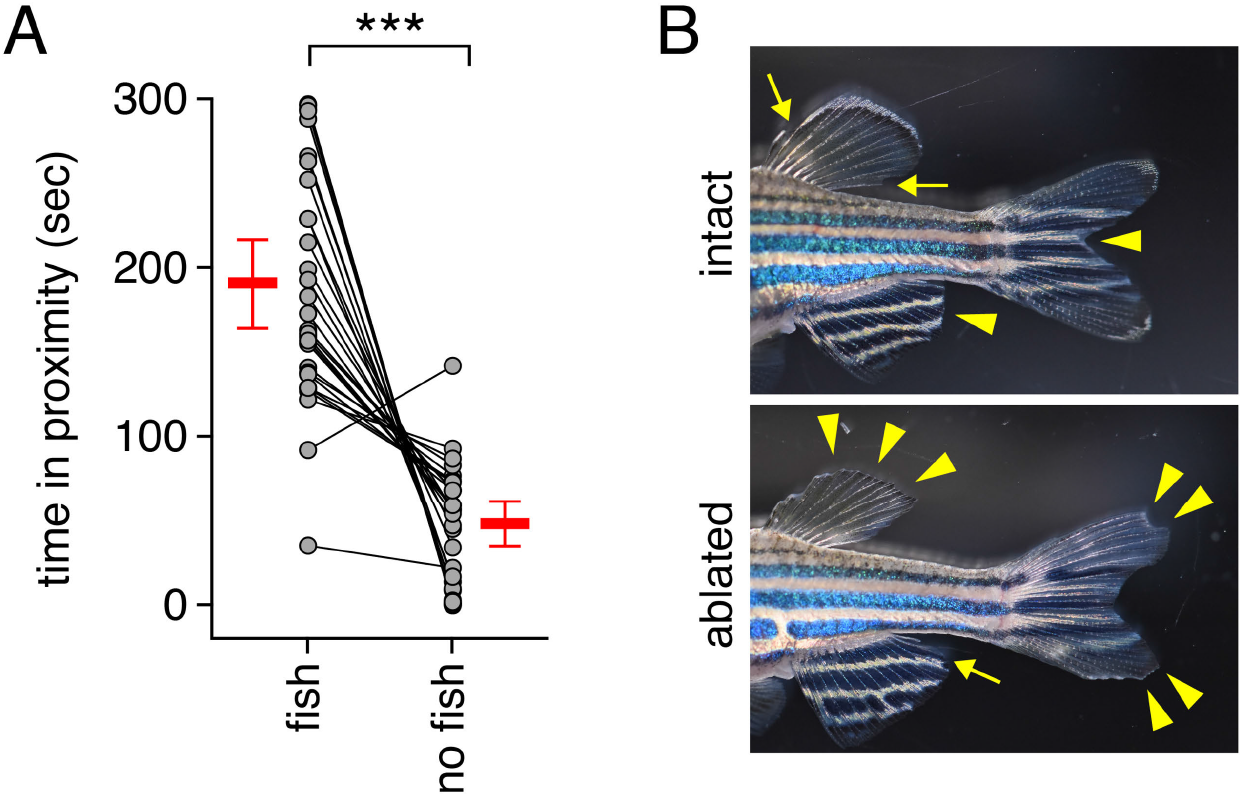
Behavioral responses to melanoleucophore complement. **(A)** Preliminary experiments confirmed suitability of test arena, as fish spent more time in association with a stimulus compartment containing a shoal of fish than a stimulus compartment without fish (difference in time spent between fish and no-fish compartments, null hypothesis mean = 0 sec: two-tailed, *t*_28_=7.71, *P*<0.0001). Each set of points connected by a line indicates times spent by an individual fish in proximity to one or another stimulus shoal compartment and bars indicate means ± 95% CI. Analyses of variance did not reveal differences in times spent shoaling associated with sex of test fish, shoaling fish, or interactions between the two (all *P*>0.4). **(B)** Examples of stimulus fish as presented to test fish in behavioral trials, following sham-manipulation (intact) or excision of melanoleucophore-containing fin tissue (ablated) two days prior to testing. Arrows indicate locations of cuts without tissue removal whereas arrowheads indicate regions from which tissue was removed (see Materials & Methods).

**Fig. S9.**
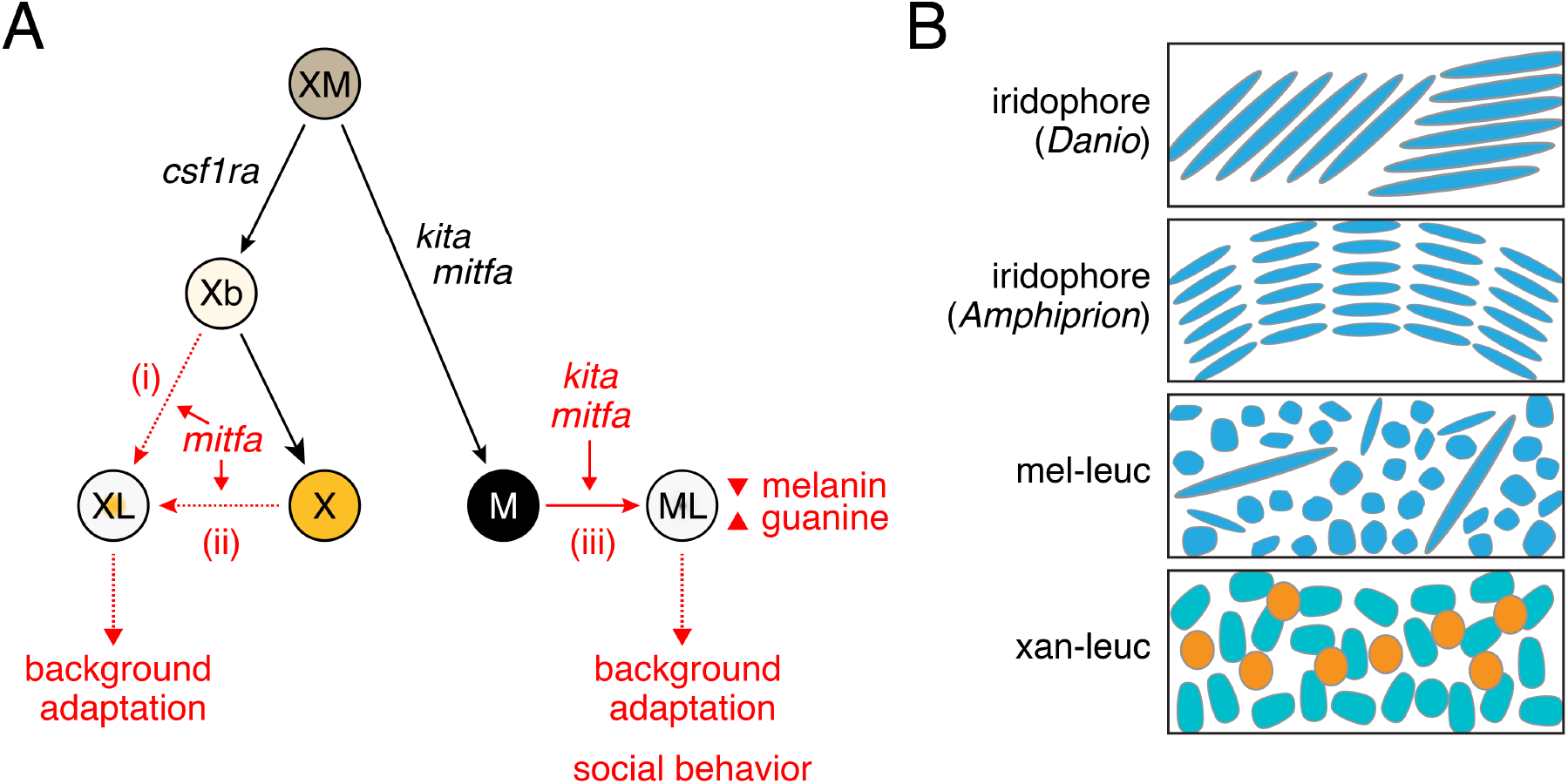
Summary of observations. **(A)** Integrated results from this study (red) and prior studies of the caudal fin [black; (13, 14)]. Dotted lines are hypothetical. *Danio* xantholeucophores (XL) and melanoleucophores (ML) arise from different sublineages, likely having a common progenitor (XM) that gives rises to xanthophores (X) and melanophores (M). Our results do not distinguish between either of two possible progenitors of xantholeucophores, which could arise directly from unpigmented xanthoblasts (Xb; i), or by conversion of differentiated xanthophores (X; ii); melanoleucophores, however, arise directly from melanophores (iii). Each leucophore lineage is constrained by prior lineage requirements (*csf1ra; kita, mitfa*), yet melanoleucophore and xantholeucophore specific roles for *mitfa* were identified, as was a melanoleucophore-specific role for *kita*. Differentiation of melanoleucophores further involved reduced expression of melanin pathway genes and loss of melanin by still unknown mechanisms, and the simultaneous upregulation of *de novo* purine synthesis genes, with deposition of crystalline guanine. Both leucophores responded in the context of background adaptation, and melanoleucophore complement was associated with differences in approach and shoaling behavior, raising the possibility of ecological roles in nature. **(B)** Pigment organelle morphologies and arrangements in light-reflecting pigment cells suggest parallel evolution of white pigment phenotype. Stacked reflecting platelets of iridophores (e.g., *Danio)* are responsible for iridescence, whereas stacked, but radially arranged platelets [*Amphiprion;* (15)] are associated with a white, rather than iridescent phenotype. By contrast, presumptive guanine-containing organelles of melanoleucophores are irregularly arranged, as are pterinosomes (green) that contain pteridines and the presumptive white pigment of xantholeucophores (orange, carotenoid vesicles).

## Materials and Methods

### Fish Stocks, Staging, Rearing Conditions and Handling

Staging used standardized standard length [SSL; (6)] and fish were at late juvenile or adult stages (>14 SSL) except as indicated. Fish were maintained at ~28.5C with 10 h : 14 h light:dark conditions. Larvae were reared with marine rotifers supplemented with Artemac (Aquafauna) followed by Artemia and flake food. Adults were maintained on Artemia and flake food. Fish stocks of *Danio rerio:* WT(ABb), a derivative of inbred AB^wp^, *Tg(tyrp1b:palm-mCherry)^wp.rt11^* (16) *Tg(aox5:palmEGFP)^wp.rt22^*(17); *Tg(pnp4a:palmmCherry)^wp.rt10^* (17); *kita^b5^* (8) *kita^ji199^* (18) *csf1ra^j4blue^*(7) *ltk^j9s1^* (9) *mitfa^w2^* (10) *mitfa^vc7^* (19) *scarb1^vp.r32c1^, tyr^vp.r33c1^* (5); *slc45a2^b4^* (20). Wild-type *Danio* species other than *D. rerio* were field-collected or obtained from commercial suppliers and maintained in the laboratory for 1–10 generations. Fish were anesthetized in MS222 prior to clipping fins or imaging. To contract pigment granules for imaging, fish were treated with 1 mg / ml epinephrine for ~5 min. All procedures involving live fish followed federal, state and local guidelines for humane treatment and protocols approved by Institutional Animal Care and Use Committees of University of Virginia and University of Washington.

### CRISPR/Cas9 Mutagenesis

One-cell embryos were injected with 200 ng/μl sgRNAs and 500 ng/μl Cas9 protein (PNA Bio) according to standard procedures (21). Guides were tested for mutagenicity by Sanger sequencing and injected fish were reared to maturity at which time they were screened for germline transmission of new mutations. Mutants of *D. aesculapii* generated for studies of adult body pigmentation (EJB and DMP, manuscript in preparation) were F1–F3 progeny of injected fish and heteroallelic for loss of function mutations resembling null alleles of *D. rerio*. sgRNA target sites, excluding PAM were: *csf1ra*, GGATCAGGACACCCTTTCTG, GGTTGTAAAACCTGTTCTCT; *kita*, GGGAAAATATTCATGCCGAG, GGACCTTGTGGGGTAATGGT; *mitfa*, GGGAGGGTGAGAAGGGCCAT, GGGCTCCCAGTGCACTGGAC.

### Transgenesis and Lineage Analysis

Tol2 mediated transgenesis followed established methods (22). A *mitfa:tdTomato* construct was generated by Gateway cloning using the Tol2Kit (23) using a 5 kb upstream regulatory region of *mitfa* and *tdTomato*. A *pnp4a:nlsEos* construct was generated by ligation of 9.4 kb *pnp4a* regulatory sequence to nuclear localizing, multimeric EosFP fluorophore. For mosaic injections in clonal labeling and Eos fate mapping, 5 pg plasmid DNA and 20 pg Tol2 mRNA were injected into 1–2 cell embryos (14, 24). For clonal labeling, 494 injected embryos were screened at 5 dpf for tdTomato+ melanophores, yielding 228 (46% of injected) larvae that were reared to 12 SSL before screening again for tdTomato in the dorsal fin, resulting in 64 (13% of injected) individuals, which were then scored for labeled melanophores and melanoleucophores. The infrequent labeling of dorsal fins suggested we had labeled only single clonal lineages and the distributions of these cells, limited to regions of 1–2 fins and extending proximal–distal along the fins, were consistent with prior analyses of clonally related fin pigment cells (13, 14); in only a single individual were labeled pigment cells non-contiguous, occurring at fin rays 2–3 and 5–6, and this fish was excluded from analysis. For Eos-labeling, embryos were screened at 5 dpf for *pnp4a*:Eos+ iridophores then reared in tanks shaded from ambient light to prevent spontaneous photoconversion (17). Eos+ fish were screened again at 8.6 SSL for labeling in the dorsal fin, at which time Eos+ melanophore clones were photoconverted (i.e., green to red).

### Imaging

Images were acquired with a Zeiss AxioObserver epifluorescence inverted microscope, a Zeiss LSM 880 laser scanning confocal microscope equipped with Fast Airyscan detector, a AxioZoom V.16 stereomicroscope using Axiocam 506 color cameras, and a Leica M205FA stereomicroscope with Leica DFC550 color camera. Images were adjusted for display level, corrected for color balance, and aligned if necessary to correct for imaging on different platforms in Adobe Photoshop CS. Images from all conditions within each analysis were treated identically.

### Image Series and Cell Counts

Fish for daily image series or cell counts were reared individually in glass beakers, treated with epinephrine to contract pigment, anesthetized using MS222, imaged and then allowed to recover. For analyses of dorsal fin melanoleucophores containing melanin, cells were counted from an image series of whole fins. For experiments using *mitfa^vc7^* and *kita^j1e99^* temperature-sensitive alleles, melanophores and melanoleucophores were counted manually under incident light.

### Temperature Shift Experiments

Temperature shift experiments were conducted using glass tanks either heated with 300V submersible heaters or cooled with a 0.25 HP drop-in chiller. Temperature was regulated using a digital temperature regulator set to maintain ±0.5°C and cross-referenced daily with a second digital thermometer. For temperature-shift experiments using the *mitfa^vc7^* allele, fish were reared at permissive temperature (25 °C) until melanophores and melanoleucophores had fully populated the distal dorsal fin (9.2 SSL) (19). Melanophores and melanoleucophores were counted and fish were then placed in beakers and shifted to a restrictive temperature (33 °C) for 2 d at which time melanophores and melanoleucophores were counted again. For temperature-shift experiments using the *kita^j1e99^* allele, fish were reared at permissive temperature (25 °C), intermediate temperature (28 °C), or restrictive temperature (33 °C) until melanophores and melanoleucophores of controls at permissive temperature had populated the distal dorsal fin (9.2 SSL) (18, 25).

### Transmission Electron Microscopy

Fin portions were amputated and prefixed in 1:1 4% glutaraldehyde and PBS at room temperature for 30 minutes to maintain structure. Fins were then fixed in sodium cacodylate buffered 4% glutaraldehyde overnight at 4°C. Tissue was post-fixed in 2% osmium tetroxide, block stained in 1% uranyl acetate overnight at 4°C, then dehydrated with an ethanol series. Tissue was then infiltrated with a 1:1 propylene oxide:Durcupan resin for 2 hours followed by fresh Durcupan resin overnight and flat embedded prior to polymerization. Blocks were thin sectioned on a Leica EM UC7 and sections images on a JEOL 1230 transmission electron microscope.

### Hyperspectral Imaging

Amputated fins were placed in PBS at room temperature and spectral measurements were performed using a PARISS hyperspectral imaging system (Lightform, Inc.), which acquires 380–980 nm spectra instantaneously from each pixel (1.25 μm x 1.25 μm) in a line of pixels. The microscope stage was then set to move adjacently to acquire another line of pixels. The PARISS software stitches these lines of pixels together to make a grayscale 2D image (reflecting the intensity of spectra) which was used to choose the spatial location of the spectra of interest. The hyperspectral imager was mounted on a Nikon Eclipse 80i microscope. Spectral calibration was performed using a MIDL Hg^+^/Ar^+^ wavelength calibration lamp (Lightform Inc.) with accuracy better than 2 nm. Cells spectra were acquired under both specular reflectance and transmittance mode, using a tungsten halogen light source with a Nikon NCB11 filter, and a 20x 0.50 NA objective. Reflectance spectra were normalized using a standard silver mirror (Thorlabs, Inc.). All spectra were smoothed with a moving average of 3 as conventionally performed, and plotted using Matlab.

### Pteridine Autofluorescence

To assess pteridine content of leucophores, amputed fins were imaged before and after exposure to dilute ammonia (pH 10.0), which liberates pteridines from protein carriers resulting in autoflourescence under DAPI illunination (3, 4).

### Mass Spectrometry

Each of three fin regions were excised from adult fish that were either wild-type, or mutant for *mitfa* or *ltk* (*n*=20 per genotype): (i) dorsal fin, distal region containing melanoleucophore stripe, including xanthophores as well; (ii) dorsal fin, adjacent to melanoleucophore stripe, containing only melanophores and xanthophores, and of size equivalent to i; (iii) anal fin, proximal light “interstripes” containing xantholeucophores and iridophores, as well as intervening dark stripe of melanophores. Purines were extracted from fin tissue in 50 μL 1M NaOH at 37 °C. Liquid chromatography/mass spectrometry (LCMS) was performed using an Agilent 1100 HPLC system equipped with a Waters XTerra MS C18, 5 μm, 4.6 Å 50 mm column or Poroshell 120 EC-C18 4.6 Å x 100 mm column. An Agilent photodiode array detector and an in-line Agilent 6130 single quadrupole mass spectrometer were used for detection. The analytical HPLC method used gradient elution from 0% to 95% acetonitrile in water (0.1% formic acid) over 6 min and Agilent ChemStation software were used for quantification. Chromatograms of the compounds were monitored at 254 nm, 210 nm, and 280 nm.

### Raman Spectroscopy

Fin portions were amputated and placed in PBS. Guanine references were prepared by recrystallizing guanine powder (Sigma Aldrich, G6779) (26). Samples were prepared by sandwiching prepared tissues or crystals between quartz coverslips (Ted Pella, Inc., Cat. No. 26016) and glass microscope slides (VWR, Cat. No. 16004-422). All data were collected at RT using a custom-built Raman microscope (27). Briefly, the 514-nm line of an argon-ion laser (CVI Melles Griot, 35-MAP-431-200) was passed through a clean-up filter (Semrock, LL01-514-25) and then directed into a modified inverted microscope (Olympus IX71). Excitation light (~30 mW at the sample) was directed to the sample using a dichroic mirror (Semrock, LPD01-514RU-25x36-1.1) and a 60x long-working distance objective (Olympus, LUCPlanFL N 60X/0.70). Spontaneous Raman Stokes scattering was collected through the same objective, filtered (Semrock, LP02-514RE-25) to remove any residual excitation light or Rayleigh scattering, and then directed into a 320 mm focal length (f/4.1 aperture) imaging spectrometer (Horiba Scientific, iHR 320) first through a 400 μm pinhole (Thorlabs) and a 50 μm slit, and dispersed using a 1200 g/mm grating. Individual spectra were collected for 5–7 s acquisitions (x25–30) from 500–3700 cm^−1^ with high gain enabled on a liquid nitrogen cooled, back illuminated deep-depletion CCD array (Horiba Scientific, Symphony II, 1024 × 256 px, 26.6 mm × 6.6 mm, 1 MHz repetition rate). Bright field images were collected using a USB 2.0 camera (iDS, UI-1220-C). Daily calibration of imaging spectrometer was done using neat cyclohexane (20 μL in a sealed capillary tube). Bandpass and accuracy were found to be <12 cm^−1^ and ±1 cm^−1^, respectively. All Raman spectra were corrected by applying a baseline polynomial fit (Lab Spec 6 software).

### Isolation of Cells for Single Cell RNA-Sequencing

Fins were amputated from fish expressing both *pnp4a:palmmCherry* and *tyrp1b:palmmCherry*. To normalize capture of relevant cell types distal dorsal fin regions (~10 mm standard length, SL; *n*=20) and proximal interstripe anal fin regions (~14 mm SL; *n*=10) were extracted. Tissue was enzymatically dissociated with Liberase (0.25 mg/mL in dPBS) at 25°C for 15 min followed by manual trituration with a flame polished glass pipette for 5 min. Cell suspensions were then filtered through a 70 μm Nylon cell strainer to obtain a single cell suspension. Liberated cells were re-suspended in 1% BSA / 5% FBS in dPBS and DAPI (0.1 μg/mL, 15 min) before FACS purification. All plastic and glass surfaces of cell contact were coated with 1% BSA in dPBS before to use. Prior to sorting for fluorescence levels, single cells were isolated by sequentially gating cells according to their SSC-A vs. FSC-A, FSC-H vs FSC-W and SSC-H vs SSC-W profiles according to standard flow cytometry practices. Cells with high levels of DAPI staining were excluded as dead or damaged. Cells from wild-type zebrafish were used as negative control to determine gates for detection of mCherry and GFP fluorescence, and then cells from transgenic fish were purified according to these gates. All samples were kept on ice, except during Liberase incubation, and then sorted chilled.

### Single Cell Collection, Library Construction and Sequencing

Following FACS, cells were pelleted and resuspended in 0.04% ultrapure BSA (ThermoFisher Scientific). We targeted 4000 cells for capture in a single lane using the Chromium platform (10X Genomics). Single-cell mRNA libraries were prepared using the single-cell 3’ solution V2 kit (10X Genomics). Quality control and quantification assays were performed using a Qubit fluorometer (Thermo Fisher) and a D1000 Screentape Assay (Agilent). Libraries were sequenced on an Illumina NextSeq 500 using 75-cycle, high output kits (read 1: 26 cycles, i7 Index: 8 cycles, read 2: 57 cycles). The full library was sequenced to an average depth of ~130 million reads. This resulted in an average read depth of 28,000 reads/cell.

### scRNA-Seq Data Processing

We built a zebrafish STAR genome index using gene annotations from Ensembl GRCz10 plus manually annotated entries for mCherry transcript, filtered for protein-coding genes (with Cell Ranger *mkgtf* and *mkref* options). Final cellular barcodes and Unique Molecular Identifiers (UMIs) were determined using Cell Ranger 2.0.2 (10X Genomics) and cells were filtered to include only high-quality cells. Cell Ranger defaults for selecting cell-associated barcodes versus barcodes associated with empty partitions were used, resulting in a final gene-barcode matrix containing 4,594 barcoded cells and gene expression counts.

### UMAP Visualization and Clustering

We used Uniform Manifold Approximation and Projection (UMAP) (28) to project cells in two dimensions and performed louvain clustering (29) using the reduceDimension and clusterCells functions in Monocle (v.2.99.1) using default parameters (except for, reduceDimension: reduction_method=UMAP, metric=cosine, n_neighbors=20, mid_dist=0.4; clusterCells: res=1e-3, k=25). We assigned clusters to cell types based on comparison of genes detected (Table 1) to published cell-type specific markers. All genes were given as input to Principal Components Analysis (PCA). The top 30 principal components (high-loading, based on the associated scree plot) were then used as input to UMAP for generating either 2D projections of the data. For subclustering of pigment cell clusters (melanophores, melanoleucophores, xanthophores, and xantholeucophores) we re-subsetted the data and applied UMAP dimensionality reduction and louvain clustering.

### Differential Expression Analysis to Determine Cell-type Markers

To identify cell-type specific genes, we used the principalGraphTest function in Monocle3 (v.2.99.1) with default parameters (30). This function uses a spatial correlation analysis, the Moran’s I test, to assess spatially restricted gene expression patterns in low dimensional space. We selected markers by optimizing for high specificity, expression levels and effect sizes within clusters (Table 2). Expression levels presented are normalized by size factor (ratio of total UMI counts for each cell and geometric mean of total UMI from all cells) (30).

### Pigment Cell Subcluster Assignment and Verification

Pigment cell subclusters were broadly assigned as either xanthophores and xantholeucophores or melanophores and melanoleucophores based on *aox5* or *tyrp1b* expression (Fig. S6F). To further distinguish among these cell types we used RT-PCR to assay the top marker identified by scRNA-Seq for each cell type by specificity and expression (Table 2). Distal dorsal fin tissue containing melanoleucophores, melanophores and xanthophores, or tissue from the proximal two interstripe regions of the anal fin, containing xantholeucophores, was dissociated (see above) and 100 cells of each type collected by mouth-pipette using a needle that had been coated in sterile 5% FBS / 1% BSA overnight at 4 °C. Cells were identified by phenotype under incident light and washed (expelled and recollected) 3 times in dPBS before final collection in lysis buffer for mRNA capture. mRNA was collected using the Ambion RNAqueous-Micro kit and cDNA was synthesized using oligo-dT primed SuperScript IV reverse transcriptase. Primers were designed to have at least one sequence that overlapped and exon/intron boundary. PCR using Q5 High-Fidelity DNA Polymerase ran for 30 cycles at 98 °C—10 sec, 60 °C—20 sec, 72 °C—30 sec and amplicons were separated on a 1% agarose gel. Primers were: entpd3(f)-cttatgtttcccgtggactaaagta, entpd3(r)-attggatctcccttcatttgtcata; ocaa2(f)-tattgtcgttctgtgttcactgttt, oca2(r)-attatacagcacctgctgagttctt; sfrp1a(f)-gtgagtttgccattaagaccaagat, sfrp1a(r)-gagaaggtactgtttgtccacctt; scinla(f)-ggtaatgagtgttctcaggatgaaa, scinla(r)-attgatagatgtccttgccaagat.

### Trajectory Analysis

The top 100 highly dispersed genes within melanophore and melanoleucophore clusters were chosen as feature genes to resolve pseudotemporal trajectories using the setOrderingFilter, reduceDimension, and orderCells functions in Monocle (v2.9.0) using default parameters with the exception of setting max_components = 2 and num_dim = 20 to generate the trajectory in 2D with the top 20 PCs (high-loading based on scree plot) during dimensionality reduction.

### Development and Analysis of Pathway Signature Scores

Gene sets for signature scores for purine synthesis (31), melanin synthesis (32) pteridine synthesis (3) and carotenoid deposition (33) were manually curated from the literature and ZFIN (34). Signature scores were calculated by generating z-scores [using scale()] of the mean of expression values (log transformed, size factor normalized) from genes in a given set.

### Physiological Response Testing

To evaluate physiological response during background adaptation to light or dark backgrounds individual fish (total *n*=6; *n*=3 per replicate with two replicates per background) were adapted in a 500 mL black (dark background) or white (light background) colored beaker under constant light for 5 minutes. Fish were then anesthetized with MS222 in the test beaker and cells from a central inter-ray region of the distal dorsal fin (melanophores and melanoleucophores) or of the anal fin second interstripe (proximal-distal; xantholeucophores) were scored under incident light. Cells were scored as either contracted or dispersed according to stages 1–3 or 4–5, respectively, of the Hogben and Slome scale (35).

### Behavioral Assays

#### Testing Arena

To test for a behavioral response to the presence of melanoleucophores, wild-type NHGR1 strain *D. rerio* were chosen for the uniformity of their pigment cell complements and limited allelic variation (36). Fish were subjected to assays consisting of three location options: open water (i.e., center third of tank), or proximity to conspecifics on either end of the tank (left or right thirds). The tank (30 cm long, 14 cm wide x 18 cm tall) was divided into three chambers with Plexiglass transparent to UV: a test chamber in the central portion of the tank (18 cm long x 14 cm wide x 18 cm tall) and two chambers for stimulus shoals on either side (each 6 cm long x 14 cm wide x 18 cm tall). For testing, chambers were filled with system water to a depth of 14 cm. Full spectrum LED lights (Draco Broadcast Dracast LED500, 5600K) were positioned 1 m above the tank to allow even lighting (~5500 lux) across the entire test arena. To avoid disturbance, the tank was placed in an open-top black box 52 cm long x 40 cm wide x 48 cm tall) with a hole 24 cm long x 22 cm tall) for a video camera cut in one side. Black plastic was draped around the camera to prevent fish from viewing the environment beyond the assay arena. All fish were recorded at 30 frames per sec with a Panasonic HC-V770 digital camera.

#### First Approach and Shoaling Assays

Stimulus shoals comprising two fish of the same sex were placed into the tank using a fish net and acclimated in the shoal chambers for 10 min prior to the onset of each testing round. Each testing round consisted of four test fish (10 min per fish; 40 min total). Test fish were dropped into the acclimation chamber (opaque cylinder, 4 cm inner diameter) using a fish net. After 5 min of acclimation, test fish were released into the test chamber for 5 min. After assaying the behavior of each test fish and after a complete round of testing, test fish and shoal fish were removed from the tank using a fish net. Between each testing round, the tank was rinsed with 65 °C water for 10 min followed by a brief rinse with fish system water before all chambers of the tank were refilled with system water. To control for possible circadian effects on sensory perception, motivation, or locomotion, all fish were assayed between 8 and 12 h after lights-on (13:00 and 17:00).

In preliminary assays, the suitability of the testing arena was confirmed by verifying that test fish would preferentially shoal with other fish as expected (37, 38), when presented with compartments either containing fish or not containing fish (Fig. S8*B*). Preference towards fish or empty chamber was tested with a reciprocal design. Male test fish (M; *n*=15 total) were tested with either a male (m; *n*=8 tests) or female (f; *n*=7 tests) shoal. Likewise, female test fish (F; *n*=14) were tested with either a male (m; *n*=7 tests) or female (f; *n*=7 tests) shoal. Placement of the shoal was randomized for each round of testing. Test fish were scored for total times spent in each of three segments of the test chamber: the 6 cm closest to the stimulus shoal, the 6 cm closest to the empty compartment, or the 6 cm in the central portion of the test chamber. For testing the preference of test fish to shoal in the vicinity of stimulus fish vs. an empty compartment, we calculated seconds spent swimming within 6 cm of the stimulus shoal – seconds spent swimming within 6 cm of the empty stimulus compartment; under a null hypothesis of no preference the mean for this value should be 0 sec.

To test for individual abilities to detect and respond to melanoleucophores, fish either had tissue containing melanoleucophores removed (ablated) or had melanoleucophore containing tissue intact (sham manipulated). Each melanoleucophore-ablated fish had a thin strip of melanoleucophore-containing tissue removed from the distal edge of the dorsal fin and the distal tips of the caudal fin with microdissection scissors and fine forceps (Fig. S8*B*). Sham-manipulated fish were lesioned with single inward cuts to the dorsal fin, without tissue removal, at anterior and posterior edges. To control for effects of tissue removal in general, sham-manipulated fish also had tissue excised from the central portion of the caudal fin, between the dorsal and ventral lobes, and along the caudal most edge of the anal fin. Melanoleucophore-ablated fish were correspondingly lesioned with a single inward cut on the rostral edge of the area removed from the anal fin of sham-manipulated fish. Prior to testing, all fish were allowed to recover and partially regenerate lost tissue, though not melanoleucophores, for 2 d in home tanks.

Ablated fish were housed with other ablated fish whereas sham-manipulated fish were housed with other sham-manipulated fish, with adjacent tanks alternating between lesion type so that all fish were in visual proximity to both ablated and sham-manipulated fish. Because stimulus shoals might respond differently to test fish with or without melanoleucophores, and such effects might influence test fish responses, both test fish and stimulus fish were assigned randomly to melanoleucophore-ablated or sham-manipulated groups. Preliminary analyses did not reveal significant effects on test fish behavior depending on whether test fish themselves were melanoleucophore-ablated or sham-manipulated (*P*>0.5); this factor was removed from subsequent analyses.

For testing effects of melanoleucophore complement on behavior, male test fish (M; *n*=18) were presented with two shoals of the same sex, one of which was melanoleucophore-ablated and one of which was sham-manipulated (male stimulus shoals, m: *n*=9 tests; female stimulus shoals, f: *n*=7 tests). Female test fish (F; *n*=17) were likewise presented with same sex melanoleucophore-ablated or sham-manipulated same-sex shoals (male stimulus shoals, m: *n*=7 tests; female stimulus shoals, f: *n*=10 tests). Placement side of ablated vs. sham-manipulated stimulus shoals was randomized for each round of testing; for simplicity, sides containing sham-manipulated fish and melanoleucophore-ablated fish are depicted as left and right, respectively, in Fig. 4*C* and Fig. 8C. Test fish were assayed for total time shoaling in proximity to ablated vs. intact stimulus fish according to time accumulated in each third of the test chamber closest to one shoal or the other. Test fish were also assayed to determine whether ablated or sham-manipulated fish were first approached upon release into the test chamber.

#### Behavior Quantification

Videos of fish during the acclimation and test phases were scored using Tracker Video Analysis and Modeling Tool software (version 5.0.5; https://physlets.org/tracker). Frames per step was set to 30 to score test fish position every 1 sec. Fish position was scored for the entire duration of the shoaling test (i.e., 5 min or 300 steps) using the Point Mass feature with mass set to the default (1.0 kg). The coordinate system was established with the point of origin at the shoal (fish vs. no fish assay) or the melanoleucophore-ablated fish (melanoleucophore response assay). Parameters quantified included x position and y position. The test arena was divided evenly into three domains along the x-axis. The position of first approach was defined as the closest x position of the first approach within either of the two domains closest to the shoal chambers.

### Statistical analysis

Analyses were performed using JMP 14.0 for Macintosh (SAS Institute, Cary NC) or *R* [version 3.5.0] (39)

